# Spatiotemporal pectin remodelling, glycoproteins, and LEA proteins maintain cell wall integrity during desiccation and rehydration in *Ramonda serbica*

**DOI:** 10.64898/2026.09.03.749093

**Authors:** Ana Pantelic, Tatiana Ilina, Dejana Milić, Nataliia Kutyrieva-Nowak, Stefan Lekić, Lazar Popović, Thierry Balliau, Ljubodrag Vujisić, Agata Leszczuk, Mélisande Blein-Nicolas, Marija Vidović

## Abstract

Vegetative desiccation tolerance requires specialised cell wall (CW) adaptations to withstand severe mechanical stress during dehydration and rehydration. While intracellular protective strategies in resurrection plants, including *Ramonda serbica*, are well documented, the CW response remains poorly understood. Here, we integrated immunocytochemical profiling, FTIR spectroscopy, quantification of CW-bound phenolics, transcriptomics, and ionically bound CW proteomics across hydrated (HL), desiccated (DL), and rehydrated states (R1–1h, R2–24h, R3–48h).

Reversible CW folding was facilitated by condensed arabinogalactan-proteins (AGPs) and extensins, alongside a site-specific balance between pectin methylesterification and demethylesterification. Structural compaction was further reinforced by the accumulation of CW-bound hydroxycinnamates, which persisted through R1 phase. Moreover, basic 7S globulin, miraculin, α-galactosidase, and two LEA4 protein family members were strongly accumulated during DL and R1, providing the first evidence of ionically CW-bound LEA proteins. Initial rewatering (R1) triggered a rapid transcriptomic reactivation of pectin-degrading/modifying enzymes, carbohydrate-active enzymes, and subtilases, accompanied by unesterified pectin enrichment. By 24–48 h (R2–R3), CW-bound hydroxycinnamic acids declined, and CW architecture, gene expression, and proteome profiles returned to baseline levels. Overall, our findings reveal a coordinated spatiotemporal apoplastic network—driven by glycoproteins, pectin modulation, CW-bound hydroxycinnamates, and LEA4 proteins—essential for rapid desiccation recovery.

## 1. Introduction

Resurrection plants represent a unique group of vegetative desiccation-tolerant angiosperms capable of surviving extreme cellular dehydration—losing over 90% of their relative water content (RWC)—and fully restoring their metabolic, physiological, and structural functions upon rewatering (Mitra et al. 2013; Bartels et al. 2025). During leaf desiccation, severe turgor loss subjects plant cells to profound hydrostatic tension and mechanical strain (Oliver et al. 2020). In non-desiccation-tolerant species, rapid volume shrinkage leads to uncontrolled irreversible plasmolysis, where the detachment of the plasma membrane from a rigid cell wall (CW) causes severe membrane strain, cortical cytoskeleton disruption, and ultimately lethal cellular collapse (Lang et al. 2014). In contrast, resurrection species undergo a tightly controlled, highly reversible folding of their mesophyll CWs, maintaining membrane–CW integrity throughout dehydration and rehydration (Vicré et al. 2004). Consistently, inward leaf rolling, a key morphological feature of desiccation tolerance in resurrection species such as *Ramonda serbica* Panč., exposes the abaxial leaf surface while keeping the palisade tissue shaded, thereby minimising light-induced photo-oxidative damage to the photosynthetic apparatus (Veljović-Jovanović et al. 2006; Rakić et al. 2015).

A central biomechanical requirement for controlled CW folding is the preservation of matrix fluidity in the dry state. During extreme drying, the polysaccharide matrix and its embedded structural proteins undergo dynamic biochemical and structural changes that largely govern the CW’s mechanical plasticity and resilience. Homogalacturonan (HG) pectins strongly influence CW compliance through their methylesterification status: unesterified HGs can bind calcium ions to form rigid, “egg-box” junction zones that promote CW stiffness, whereas methyl-esterified HGs sterically hinder cross-linking, maintaining matrix elasticity (Giarola et al. 2016; Wang et al. 2025). This dynamic tuning of pectin architecture is tightly regulated by a complex suite of pectin-modifying enzymes. Pectin methylesterases (PMEs) catalyse the demethylesterification of HG, either promoting Ca^2+^ cross-linking and wall stiffening or enabling subsequent enzymatic degradation. The fine-tuning of PME activity is mediated by pectin methylesterase inhibitors (PMEIs), which post-translationally block PME action to prevent excessive wall rigidification or premature pectin cleavage (Pecatelli et al. 2026). Concurrently, pectin-degrading enzymes such as polygalacturonases (PGs) and pectate lyases (PLs) hydrolyse or β-eliminatively cleave the polygalacturonan backbone, facilitating CW loosening and matrix reorganisation (Hocq et al. 2017). In rhamnogalacturonan I (RG-I) domains, RG lyases target the branched pectin backbone, while glycosyl hydrolases such as galactosidases and α-L-arabinofuranosidases (ARAFs) trim pectic arabinan and galactan side chains (Knoch et al. 2014). The enzymatic removal or retention of these neutral side chains directly influences pectin hydration capacity, steric hindrance, and polymer mobility.

Across various resurrection species (e.g., *Craterostigma plantagineum* and *Myrothamnus flabellifolia*), controlled CW loosening and stiffening are regulated by HG demethylesterification, arabinogalactan-protein (AGP) plasticisation, expansin activation, and class III peroxidase (POD)-mediated oxidative cross-linking (Moore et al. 2006, 2024; Chen et al. 2020). Structural HRGPs play a crucial role in maintaining cell wall integrity; extensins assemble into covalent networks, whereas highly branched AGPs physically interact with polysaccharides and pectin networks to reinforce CW architecture and preserve continuous surface organisation (Hijazi et al. 2014; Mnich et al. 2020). Hydroxyproline-rich glycoproteins (HpRGPs), including AGPs and extensins, function at the CW–plasma membrane continuum as biomechanical buffers and structural anchors (Chen et al. 2020).

While intracellular protective mechanisms—such as non-reducing sugar accumulation, antioxidant defence activation, and LEA protein induction—have been extensively documented, the mechanical and structural adaptations of the CW during severe dehydration remain less understood (Moore et al. 2013). In *R. serbica* leaves, desiccation involves a functional shift towards cyclic electron transport, downregulation of core photosystems, non-reducing sugar accumulation, and the specific induction of over 20 LEA proteins proposed to function as intracellular molecular shields and antioxidants (Pantelić et al. 2022), alongside the activation of key antioxidant enzymes, including PODs, superoxide dismutases, and polyphenol oxidases (Veljović-Jovanović et al. 2006; 2008; Rakić et al. 2015; Vidović et al. 2022). At the apoplastic level, we previously demonstrated a desiccation-induced up-regulation of genes encoding PMEs and germin-like proteins (GLPs)—both key drivers of CW restructuring and redox regulation (Vidović et al. 2022). Furthermore, we observed a 40% increase in the total content of total soluble phenolic compounds in desiccated compared with fully hydrated leaves (Vidović et al. 2022).

However, extending these studies to a CW proteome level remains particularly challenging due to the low abundance and tight matrix binding of apoplastic proteins, leaving their presence and functional coordination largely unexplored. Moreover, while desiccation responses have been the primary focus of most studies, the rapid recovery dynamics during early rehydration represent a crucial yet uncharacterised bottleneck for CW integrity. During this initial rewatering phase, cells experience sudden osmotic fluid influx, mechanical swelling, and a pronounced oxidative burst, due to fully unrecovered antioxidative systems (Sgherri et al. 1994; Veljović-Jovanović et al. 2006). Consequently, the precise spatiotemporal remodelling of the *R. serbica* CW matrix during dehydration and progressive rehydration phases remains uncharacterised. Specifically, the spatial distribution and methylesterification dynamics of AGPs and pectins, as well as the specific reorganisation of CW-bound phenolics, have yet to be directly mapped *in situ*.

In this study, we investigated the structural, immunocytochemical, proteomic, and biochemical dynamics of *R. serbica* leaf CW across five defined physiological states: fully hydrated control (HL), fully desiccated (DL), and three post-rewatering phases (R1–1 h, R2–24 h, R3–48 h). Our primary objective was to elucidate how dynamic modifications of CW components and their underlying gene expression profiles are temporally orchestrated to protect the tissue during desiccation and to withstand the profound osmotic and mechanical stress associated with rapid recovery. Deciphering how resurrection plants preserve CW integrity under extreme physical strain without undergoing lethal structural failure is fundamental to understanding vegetative desiccation tolerance.

## 2. Methods

### 2.1. Plant Material and Experimental Conditions

Mature specimens of the resurrection species *R. serbica* Panč. were collected from their natural habitat in the Sićevo Gorge near Niš (South-Eastern Serbia). The desiccation and rehydration protocol was carried out according to Veljović-Jovanović et al. (2008). Sampling was conducted across five sequential physiological stages: fully hydrated leaves (HL; baseline); fully desiccated leaves (DL, RWC below 10%); rehydration stage 1 (R1, 1 h post-rewatering); rehydration stage 2 (R2, 24 h post-rewatering); and rehydration stage 3 (R3, 48 h post-rewatering). For each physiological state, four independent biological replicates were collected (with the exception of R2, which consisted of three biological replicates). Each biological replicate represented an individual plant and comprised three pooled mature, light-exposed leaves. The harvested leaf material was immediately frozen in liquid nitrogen and stored at −80 °C until further analysis (samples for microscopy were prepared immediately upon leaf collection). RWC was measured at regular intervals during dehydration as explained in Vidović et al. (2022).

### 2.2. Immunofluorescence Labelling & Microscopy Imaging

Leaf tissue processing, fixation, resin (100% LR White) embedding, and sectioning were performed according to Kutyrieva-Nowak et al. (2025). Semi-thin sections (1 µm) were cut using an ultramicrotome equipped with a glass knife (PowerTome XL, RMC Boeckeler, USA) and mounted on poly-L-lysine-coated glass slides (Sigma, USA). To evaluate anatomical leaf adjustments, sections were stained with a 0.5% (w/v) aqueous solution of Toluidine Blue O for 30 s. Structural changes across desiccation and rehydration were observed and photographed using an Olympus BX51 microscope equipped with FluoView v. 5.0 software. Cell cross-sectional area measurements (n ≥ 30 cells per condition) were performed using ImageJ software (NIH, Bethesda, MD, USA).

Immunostaining techniques were carried out according to the protocol described in Kutyrieva-Nowak et al. (2025). Monoclonal antibodies targeting cell wall components—JIM13 and LM2 (AGPs), LM1 (extensins), LM16 (RG-I galactan), LM19 (unesterified HG), and LM20 (highly methyl-esterified HG) (Kerafast, USA)—were utilised. Imaging was conducted using an Olympus BX51 CLSM equipped with FluoView v. 5.0 software (Olympus Corporation, Japan). Primary antibodies were omitted for negative control reactions. All figures and schemes were assembled and edited using CorelDRAW X6.

### 2.3. CW Isolation, Purification and FTIR spectroscopy

CW fractions from leaves across all stages (HL, DL, R1, R2, R3) were isolated according to Vidović et al. (2022). A series of sequential extractions with organic solvents (ethanol, chloroform/methanol 1:1 v/v, and acetone) was performed to completely remove pigments, lipids, alkaloids, tannins, soluble sugars, and low-molecular-weight metabolites. The resulting purified cell wall material was dried to a constant mass and stored in a desiccator prior to structural analysis.

Fourier-transform infrared (FTIR) spectra of 19 isolated CW samples (4 of DL, 4 of HL, 4 of R1, 3 of R2, 4 of R3) were recorded using a Nicolet™ Summit FTIR spectrometer equipped with an Everest™ diamond ATR accessory (Thermo Fisher Scientific, Madison, WI, USA). For each sample, spectra were collected in the mid-infrared region (4,000–400 cm⁻¹) at a spectral resolution of 4 cm⁻¹, as an average of 64 co-added scans.

Spectral data processing and Principal Component Analysis (PCA) were executed using Quasar software v1.7.0 (Demšar et al. 2013; Toplak, et al 2021). To evaluate CW structural transitions, spectra were truncated to the statistically significant region (1,175-935 cm⁻¹). Pre-processing was performed in following order: Savitzky–Golay smoothing, second derivative, baseline correction, and Standard Normal Variate (SNV) normalisation. Unsupervised PCA was subsequently performed on the pre-processed spectral matrix to evaluate cluster separation and identify key vibrational bands contributing to CW remodelling across desiccation and rehydration.

### 2.4. Analysis of Cell-Wall-Bound Phenolic Compounds

Cell-wall-bound phenolics were liberated from purified CW pellets via alkaline hydrolysis using 1 M NaOH at 80 °C for 17 h in the dark at room temperature. After neutralisation, phenolic profiles were analysed by HPLC according to Vidović et al. (2022). Analyses were performed by HPLC coupled with a photodiode array detector (Ultimate 3000, Thermo Fisher Scientific, USA) on 250 × 4.6 mm, 5.0 mm, Luna C18 (2) reversed-phase column (Phenomenex Ltd. Torrance, CA, USA). Individual phenolics were identified by comparing their absorption spectra with those of authentic standards and confirmed by spiking. Quantification was based on the peak area, using Chromeleon CDS 6.8 software (Thermo Fisher Scientific, Sunnyvale, CA, USA).

### 2.5. Proteomic Analysis of CW-Enriched Fractions

#### 2.5.1. Extraction of Ionically Bound CW Proteins

CW proteins were extracted according to Printz et al. (2015). Leaf tissue was homogenised under liquid nitrogen and resuspended in isoosmotic extraction buffer (5 mM sodium acetate pH 4.6, 0.4 M sucrose, 20 mM ascorbate, 10 mM tris(2-carboxyethyl)phosphine, TCEP) to minimise cell rupture and organellar contamination. Following 30 min extraction on ice, samples were centrifuged (11,000 × *g*, 20 min, 4 °C). Extraction was repeated sequentially using buffer containing 0.6 M and then 1.0 M sucrose to remove residual intracellular proteins. Pellets were washed three times (30 min each) with sucrose-free extraction buffer to yield clean CW-enriched fractions.

Ionically bound CW proteins were sequentially extracted using differential salt solutions at 4 °C. Each extraction step was followed by centrifugation at 1,000 × *g* for 15 min at 4 °C to collect the respective supernatants: fraction 1 (weakly bound): two 30 min extractions with 200 mM CaCl2 in starting buffer; fraction 2 (pectin-associated): two 60 min extractions with 50 mM ethylene glycol tetraacetic acid (EGTA) in starting buffer; fraction 3 (strongly bound): by an overnight incubation (16 h) in 3 M LiCl at 4 °C under continuous agitation. For each sample, the supernatants from all three extraction steps were pooled to form the total ionically bound apoplastic protein fraction. The pooled supernatants, as well as the remaining cell wall pellets, were lyophilised separately and stored at −20 °C.

In parallel, the same samples were subjected to a phenol-based extraction protocol according to James et al. (2025) for comparison with the protocol for ionically bound proteins.

#### 2.5.2. Protein Digestion

Pooled ionically bound proteins (the same amount) were resolved on short-run sodium dodecyl sulphate– polyacrylamide gel electrophoresis (SDS-PAGE) gels (migration front ∼1.5 cm). Gel lanes were split into three equal parts, and the middle strip was cut into six gel cubes (∼ 1-1.5 cm^2^), on which digestion was performed according to Balliau et al. (2018). Gel pieces were dehydrated using 30 μL of 10% acetic acid / 40% ethanol. Re-hydration/washing was performed for 15 min three times with 30 μL of Solution A: 25 mM NH_4_HCO_3_ (pH 8.0) in 25% acetonitrile (ACN). Dehydration was completed with 100% ACN, followed by reduction with 50 μL of 10 mM dithiothreitol, (DTT) at 56 °C for 5 min and alkylation with 40 μL of 55 mM iodoacetamide (45 min in the dark at 37 °C). Gel pieces were washed with Solution A, shrunk with 100% ACN, and digested with 100 ng trypsin overnight at 37 °C. Supernatants were collected, combined with sequential extractions using Solution A, 0.5% trifluoroacetic acid (TFA) in 50% ACN, and pure ACN, and dried in a vacuum concentrator. Phenol-extracted proteins were digested as described in James et al. (2025).

#### 2.5.3. NanoLC-timsTOF Pro Mass Spectrometry

Peptides (100 ng per sample) were analysed using a timsTOF Pro mass spectrometer coupled to a nanoElute LC system (Bruker Daltonik, Germany). Samples (1 μL) were injected onto an Acclaim PepMap C18 trap column (0.1 ×20 mm, 5 μm, 100 Å) and separated on an Aurora C18 analytical column (75 μm ×250 mm, 1.6 μm, 120 Å; Ion Opticks, Australia) at 50 °C with a flow rate of 200 nL min^−1^. Gradient elution (Eluent A: 0.1% formic acid in water; Eluent B: 0.1% formic acid in ACN) proceeded as follows: 2-5% B (1 min), 5-13% B (18 min), 13-19% B (7 min), 19-22% B (4 min), and 95% B (7 min).

Ionisation was established via a CaptiveSpray source (1.6 kV, 180 °C, 3 L min^−1^ dry gas). Data acquisition was operated in data-dependent acquisition parallel accumulation-serial fragmentation (DDA-PASEF) mode using oTof Control (v6.0.3). The *m/z* range was set to *m/z* 100-1,700, with an ion mobility range (1/K0) of 0.7-1.1 V·s·cm⁻² K₀⁻¹ (166 ms duration; Daviere et al. (2026). Collision energy was stepped from 20 to 59 eV for charge states from 0 to 5, using an intensity threshold of 1000 counts per second (cts s⁻¹) and a target intensity of 20000 cts s⁻¹ across 6 PASEF scans within the same mobility range. The PASEF scan was repeated six times. Active exclusion was set to 24 s, resulting in a total cycle time of 1.3 seconds.

#### 2.5.4. Database Search and Bioinformatic Filtering

Peptide spectra were searched using X!Tandem Alanine (v2017.2.1.4) against a custom *R. serbica* translated transcriptomic database (Zenodo DOI: 10.5281/zenodo.6341873) and common contaminants. Parameters for the X!Tandem search are given in **Supporting Information File S1**. The mass spectrometry proteomics data have been deposited to the ProteomeXchange Consortium via the PRIDE partner repository with the dataset identifier PXD083564.

Protein inference was executed in i2MassChrQ software (v1.0.18, Langella et al. 2024) on a combined dataset of phenol-extracted and ionically-bound proteins. Proteins were selected based on an e-value of less than 10⁻5, requiring at least two unique peptides with an e-value below 0.01. Peak quantification was conducted using MassChroQ (v2.4.30; Valot et al. 2011). The mass tolerance was set to 20 ppm and quantification was performed at a level of 80% of the theoretical natural isotopic profile.

To filter out intracellular contaminants, proteins were classified as bona fide apoplastic proteins based on: (i) Signal peptide prediction by TargetP 2.0 (Almagro Armenteros et al. 2019); (ii) 70% sequence homology to proteins with TargetP-predicted signal peptides; or (iii) Prior experimental validation in the CW database (WallProtDB; Albenne et al. 2013). Due to the significant presence of intracellular contaminants among phenol-extracted proteins (data not shown), these were excluded from further processing and analysis.

#### 2.5.5. Quantitative Data Processing and Statistics

Peptide intensity data were filtered and processed in R Studio (v2024.04.2) using MCQR (v0.6.15; Balliau et al. 2025), according to the R script provided in **Supporting Information File S2**. Processing steps included intensity normalisation, knn-missing value imputation, and peptide-to-protein summarisation. Proteins were filtered using a |log2FC| ≥ 1.5 and significant differences across conditions were verified by performing a one-way ANOVA followed by a Tukey’s HSD post-hoc test (*p*_adj_ < 0.05).

### 2.6. Transcriptomic Analysis

#### 2.6.1. RNA Extraction

Total RNA was extracted according to Vidović et al. (2020). Leaf tissue (∼0.05 gDW) was homogenised in liquid nitrogen. The volume of extraction buffer was adjusted to obtain tissue-to-buffer ratios of 1:5 (w/v) for HL and R3, 1:7.5 (w/v) for R2, and 1:10 (w/v) for DL and R1. RNA concentration and purity were initially assessed using a BioSpec-nano spectrophotometer (Shimadzu Corporation, Kyoto, Japan). Prior to library preparation, RNA concentration was additionally determined using a Qubit 4.0 fluorometer with the Qubit RNA High Sensitivity Assay Kit (Thermo Fisher Scientific, Waltham, MA, USA).

#### 2.6.2. RNA Depletion, Library Preparation and Sequencing

Ribosomal RNA was depleted using the MGIEasy rRNA Depletion Kit, and sequencing libraries were prepared using the MGIEasy Fast RNA Library Prep Set according to the manufacturer’s instructions (MGI Tech Co. Ltd., Shenzhen, China). Fourteen sequencing libraries were generated from 13 biological samples representing hydrated leaves (HL, n = 3), desiccated leaves (DL, n = 3), and three successive rehydration stages (R1, n = 2; R2, n = 2; R3, n = 3). One R2 biological sample was sequenced as two technical libraries (R2 80 and R2 80N). Libraries were sequenced in paired-end mode (2 × 150 bp) on a DNBSEQ-G400 platform (MGI Tech Co. Ltd., Shenzhen, China).

The data that support the findings of this study have been deposited in Zenodo under the reserved DOI 10.5281/zenodo.22046118.

#### 2.6.3. Read Processing and De Novo Transcriptome Assembly

Raw paired-end reads were trimmed and quality-filtered using fastp (v0.23.4) Adapter sequences were detected automatically, bases with Phred quality scores below 20 were filtered, reads shorter than 50 nucleotides were removed, and paired-read overlap correction was enabled. Quality-filtered reads from all libraries were jointly assembled de novo using Trinity (v2.15.2), with a minimum transcript length of 300 nucleotides. The assembly was mapped against the published *R. serbica* transcriptome (Pantelić et al., 2022; Zenodo DOI: 10.5281/zenodo.6341873) using minimap2. Trinity transcripts were considered redundant and removed if they represented exact or near-exact matches (identity ≥ 99%, query and target coverage ≥ 98%), high-confidence equivalents (identity ≥ 97%, query and target coverage ≥ 90%), or contained/partial matches (identity ≥ 95% and query coverage ≥ 90%). Expression support was determined by preliminary Salmon mapping against the Trinity assembly and import with tximport using countsFromAbundance = “lengthScaledTPM”; counts from the two R2_80 technical libraries were summed, and transcripts with an estimated count ≥ 10 in at least two biological samples were retained. The remaining nonredundant transcripts were combined with the published transcript set to form the expanded transcript reference for downstream analyses.

#### 2.6.4. Transcript Quantification and Differential Expression Analysis

Transcript abundance was quantified using Salmon v1.10.3 with mapping validation and sequence/GC-bias corrections, then imported into R v4.4.2 via tximport v1.34.0, summing technical replicates. Differential expression was analysed with DESeq2 v1.46.0 across all ten pairwise comparisons using Wald tests. Additionally, omnibus likelihood-ratio tests (LRT) were performed for the five-condition continuum (HL vs. R1 vs. R2 vs. R3 vs. DL) and grouped states [(HL+R3) vs. R2 vs. (DL+R1)], followed by pairwise Wald post-hoc tests. Transcripts were considered differentially expressed at *p*_adj_ < 0.05 and |log2FC| ≥ 2. A conservative locus-level analysis was complementarily conducted by aggregating counts for transcripts reliably mapped to the same Pantelić gene or Trinity component.

#### 2.6.5. Functional Annotation and Selection of Cell-Wall-Related Candidates

Protein sequences were retrieved from validated translations (Pantelić et al. 2022; Zenodo DOI: 10.5281/zenodo.6340979) or where required, predicted using TransDecoder (v5.7.1, retaining only error-free coding sequences without internal stop codons or ambiguous residues. Functional annotation and domain identification were performed using DIAMOND (v2.2.5) (blastp and blastx; --very-sensitive; E-value ≤ 10⁻⁵; maximum 25 target sequences and one HSP per target; no minimum identity or query/subject coverage filters; Swiss-Prot and Gesneriaceae databases), HMMER (v3.4) (hmmscan -- cut_ga; Pfam-A), and existing eggNOG/UniProt references. N-terminal signal peptides were predicted with TargetP 2.0 (Almagro Armenteros et al. 2019), and CW-related candidates were selected by integrating TargetP outputs with apoplastic/CW functional annotations.

#### 2.6.6. Visualisation of CW-Associated Transcript Dynamics

For visualisation of the curated cell-wall-associated transcript set, variance-stabilising transformation was applied to the complete DESeq2 dataset with blind = FALSE. The 1,186 transcripts satisfying the final |log2FC| ≥ 2 selection criterion were subsequently extracted from the transformed matrix. VST expression values were averaged across biological replicates within five physiological states and standardised transcript-wise to Z-scores. Transcripts were hierarchically clustered using Euclidean distance and complete linkage, while the predefined physiological-state order (HL→DL→R1→R2→R3) was preserved by clustering rows but not columns. Heatmaps were generated in R (v4.4.2) using DESeq2 (v1.46.0) and heatmap (v1.0.13). Transcript abundances of the final CW-related transcript-level candidate set were matched by exact Transcript ID, and distinct Transcript IDs sharing the same Rs identifier were retained separately. Data were processed using Python (v3.11.15), and the resulting 100% stacked bar plots were generated in R (v4.4.2) using base graphics. DESeq2-normalised transcript counts were first averaged across biological replicates within each physiological state, after which transcript abundances were summed within each functional category.

## 3. Results

### 3.1 Tissue Architecture and Chloroplast Distribution During Desiccation and Rehydration

During desiccation (DL), leaf cells underwent a drastic reduction in cross-sectional area, shrinking to 54.5% relative to the fully hydrated state (decreasing from 5,136 µm² in HL to 2,800 µm² in DL). To accommodate this severe volume loss without mechanical rupture, cell walls exhibited extensive, accordion-like folding, accompanied by localised compaction of the CW matrix (**Figure 1, Supplementary Figure S1**).

**Figure 1.**
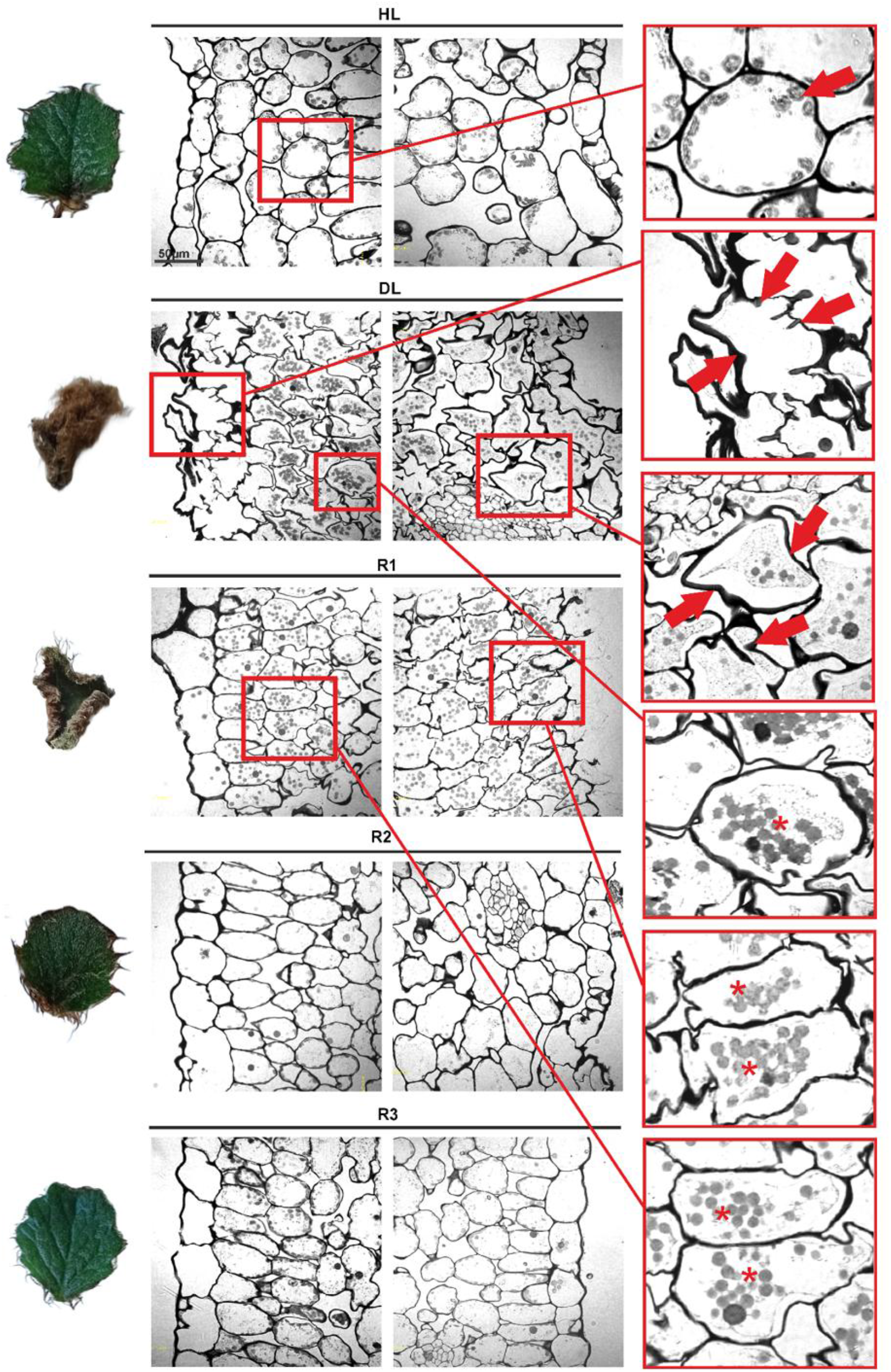
Representative light microscopy images of cross-sections of fully hydrated leaves (HL), desiccated leaves (DL), and leaves at rehydration stages (R1, R2, R3) stained with Toluidine Blue. Arrows indicate cell wall invaginations and central chloroplast clustering in DL. Asterisks (*) mark intense punctate marginal signals during R1. For each stage, the left image illustrates a cross-section showing the abaxial (lower) epidermis, while the right image illustrates a cross-section showing the adaxial (upper) leaf epidermis. Scale bars = 50 µm.

This structural response was observed across all tissue layers (Supplementary Figure S1): epidermal cells exhibited pronounced folding of their outer cell walls, rounded mesophyll cells underwent severe deformation with a marked reduction in intercellular spaces, and cells surrounding the vascular bundles— including the bundle sheath—became highly compressed. Concurrently, vacuoles were no longer observable, the protoplast underwent severe condensation towards the cell centre, and chloroplasts became densely crowded in a central aggregate.

Upon rewatering, initial water influx within one hour (R1) triggered rapid cell swelling and wall unfolding, initiating turgor recovery and partial restoration of tissue architecture. Epidermal cells regained their regular shape alongside the appearance of small internal structures (likely plastids or starch granules), mesophyll cells re-expanded with densely populated chloroplasts, and vascular tissues progressively uncompressed (**Supplementary Figure S1**). By 24 h (R2), cells recovered 95.6% of their initial HL area (reaching a mean of 4,910 µm²), accompanied by the dissipation of stress-induced vesicular signals and the repositioning of chloroplasts towards the cellular periphery.

Full histological recovery was achieved after 48 h (R3), where cells regained 95.0% of their original cross-sectional area, indicating a near-complete restoration of CW architecture towards the hydrated state, re-establishing a rounded cellular morphology with parietal, densely packed chloroplasts and stabilised cell walls (**Figure 1**). The near-complete morphological recovery across repeated desiccation–rehydration cycles suggested that no overt permanent structural damage remained in the examined tissues.

### 3.2. Immunodetection of Arabinogalactan-Proteins (AGPs), Extensins and Pectins

In hydrated leaves (HL), β-galacturonosyl-containing AGPs (JIM13) and glucuronated AGPs (LM2) were evenly distributed along the plasma membrane–CW interface (**Figure 2a**). While LM2 labelling consistently marked punctate, vesicle-like structures along the cellular periphery across all stages, the apparent signal enhancement in desiccated leaves could not be conclusively attributed to elevated vesicle density rather than a spatial concentration effect driven by protoplast shrinkage. Concurrently, extensins (LM1) formed a thin, regular network in HL that became condensed, thickened, and wrinkled during DL. This extensin scaffolding remained tightly packed around the expanding protoplast at R1 before restoring its continuous meshwork architecture at R2–R3 (**Figure 2a**).

**Figure 2.**
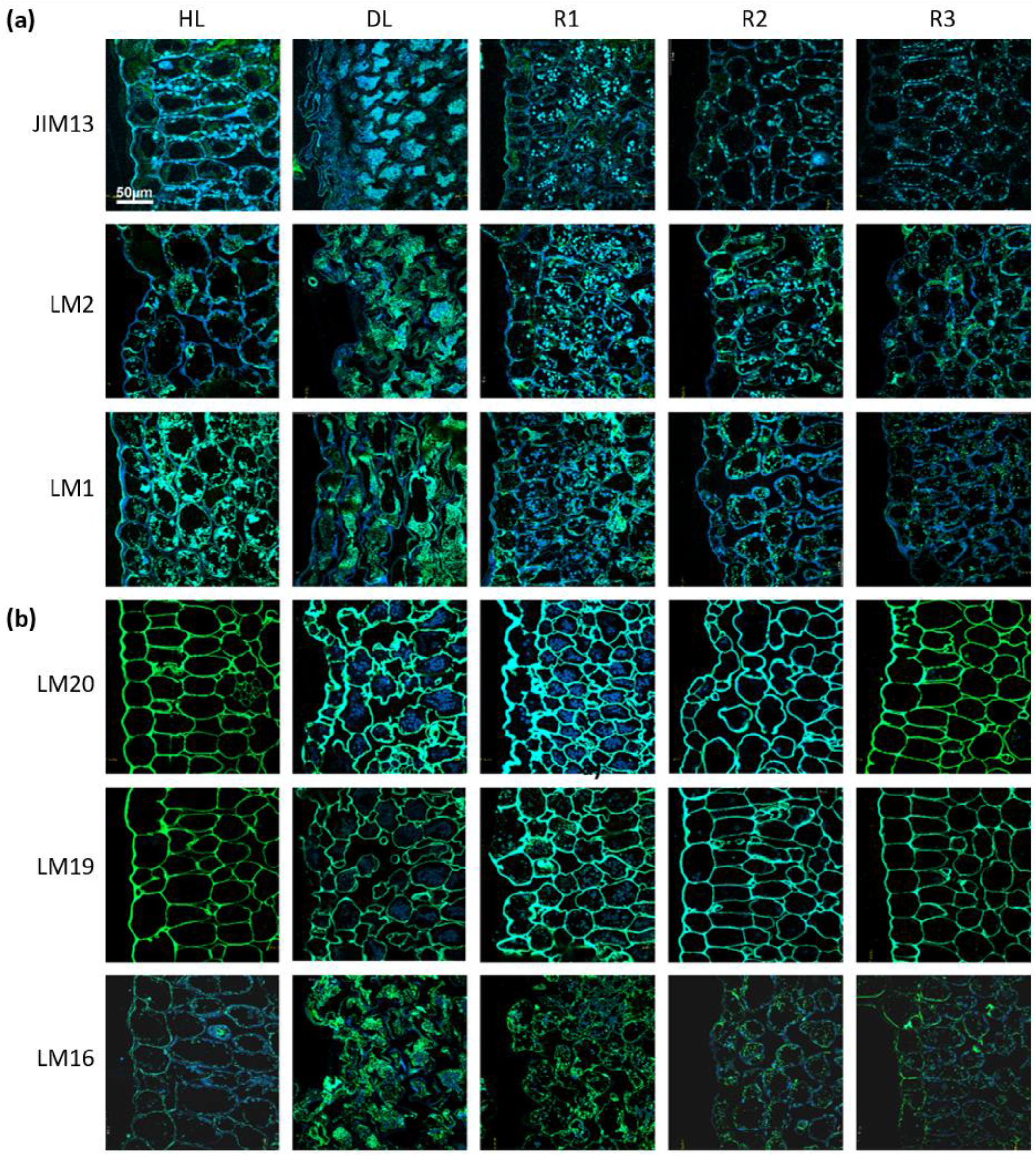
Immunofluorescent localisation of **(a)** CW glycoproteins (JIM13-recognised AGP epitopes; LM2-recognised glucuronated AGPs; LM1-recognised extensins); and **(b)** CW pectin epitopes (highly methyl-esterified HG recognised by LM20, unesterified/low methyl-esterified HG labelled by LM19, LM16-recognised RG-I galactan side-chains) across desiccation and rehydration stages (HL, DL, R1, R2, R3). Primary CWs are counterstained with Calcofluor White (blue). Scale bars = 50 µm.

Highly methyl-esterified (HG-HM; LM20), unesterified, low methyl-esterified HG (HG-LM; LM19) and homogalacturonans (HGs) co-existed in HL CWs (**Figure 2b**). In desiccated leaves, while LM20 labelling broadened and intensified across CW invaginations, LM19 epitopes were restricted to cell boundaries. Upon rehydration (R1), LM20 signal intensity dropped significantly, whereas LM19 labelling became prominent and continuous, before both restored a balanced basal distribution by R2–R3. Concurrently, LM16-recognised RG-I galactan side chains were evenly distributed in HL, showed stronger labelling across folded tissues during DL, were predominantly detected along expanding cell boundaries at R1, and showed a distribution resembling that in HL by R2–R3 (**Figure 2b**).

### 3.3. Transcriptional Profiling of CW-Related Genes

Whole-transcriptome RNA sequencing of *R. serbica* leaves across five stages (HL, DL, R1–R3) retained 268,738 transcripts after filtering, of which 42,789 were differentially expressed in at least one pairwise comparison (*p*_adj_ < 0.05 and |log2FC| ≥ 2, **Supplementary Table S1**). Overall, 87,299 differential-expression events were detected (49,288 upregulated, 38,011 downregulated).

Likelihood-ratio tests (LRT) identified 28,308 condition-dependent transcripts across five stages (HL, DL, R1–R3) and 11,447 across three merged states [(HL+R3) vs. R2 vs. (DL+R1)]. However, no transcript met the strict significance threshold across all ten pairwise comparisons or all three grouped contrasts, indicating highly stage-and contrast-dependent regulatory bursts rather than global multi-stage reconfigurations. Hierarchical clustering further illustrated stage-dependent expression patterns (**Supplementary Figure S2**).

TargetP predictions and manual functional curation identified 1186 unique differentially expressed transcripts encoding putative CW-associated proteins (|log2FC| ≥ 2, **Supplementary Table S2**). Consecutive pairwise comparisons revealed 462 DEGs in DL vs. HL, 474 in R1 vs. DL, 96 in R2 vs. R1, 2 in R3 vs. R2, and 31 in R3 vs. HL. Relative abundance of retained CW-associated DEGs categorised into seven functional groups: (1) ROS-scavenging/antioxidant enzymes, (2) stress-responsive/protective proteins, (3) pectin-modifying enzymes, (4) hemicellulose and non-pectic glycan-modifying carbohydrate-active enzymes (CAZymes), (5) structural CW proteins/glycoproteins, (6) proteases, and (7) phenolics/lignin-related proteins, alongside an “Other” group is presented in **Figure 3**.

**Figure 3.**
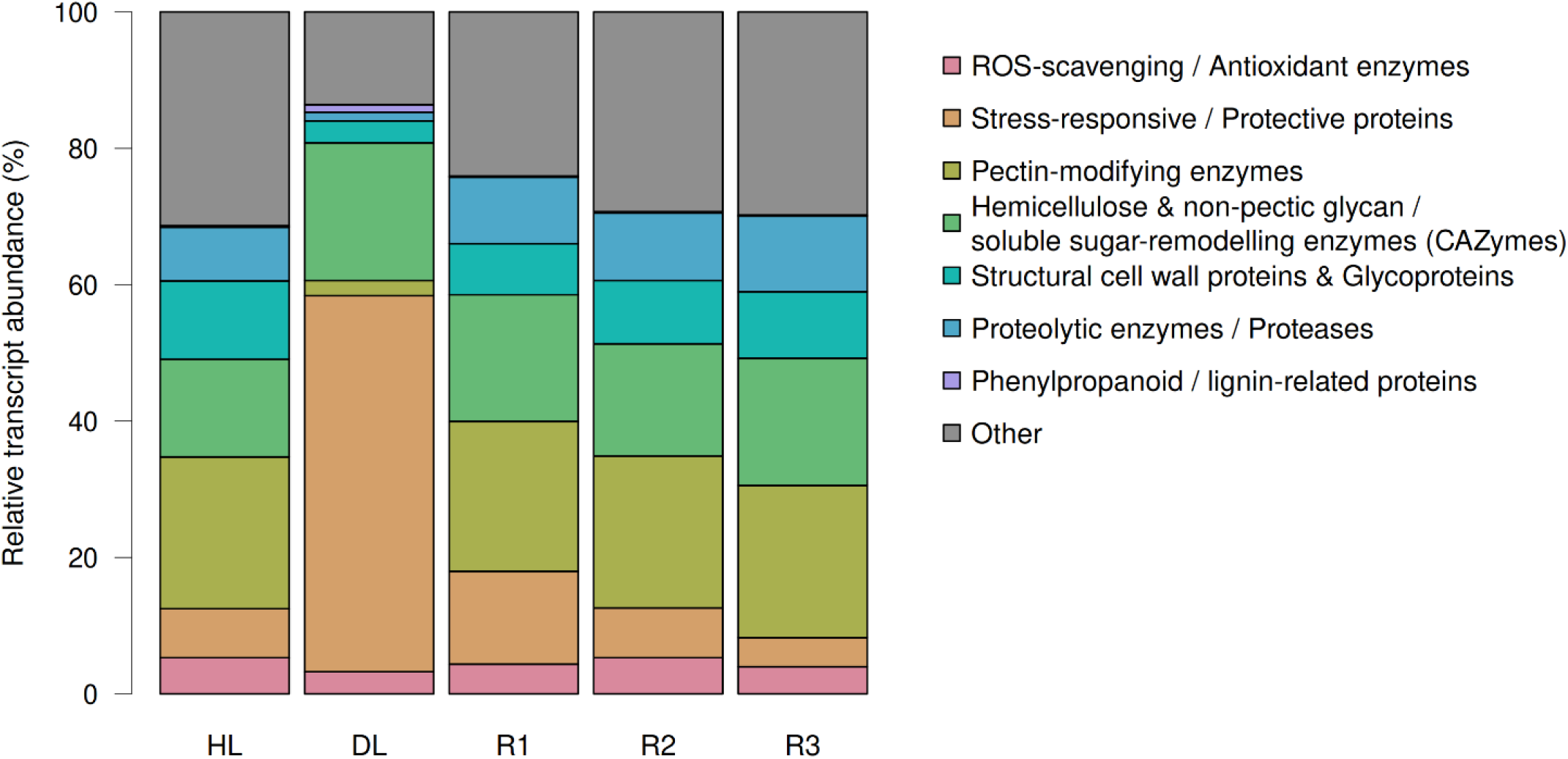
Relative abundance of CW-associated transcript categories across desiccation and rehydration. Stacked bar plot illustrating the functional distribution and composition (% of total CW-targeted transcript abundance) across five desiccation/rehydration stages: HL, DL, R1 (1 h), R2 (24 h), and R3 (48 h) **(Supplementary Table S2)**.

Desiccation induced profound transcriptomic remodelling, marked by widespread suppression of CW metabolic and structural genes alongside targeted activation of protective machinery (**Figure 3, Supplementary Figure S2, Supplementary Table S2**). DEGs downregulated in DL relative to HL were predominantly linked to CW loosening and remodelling, including CAZymes (e.g., β-glucosidases, xyloglucan endotransglucosylase/hydrolases – XTHs, β-D-xylosidases), pectin-modifying enzymes (pectin acetylesterases, PLs, PGs, and PMEs/PMEIs), structural proteins (glycine-rich proteins – GRPs, proline-rich proteins – PRPs, expansins) and key antioxidant enzymes (PODs, GLPs). Conversely, desiccation selectively upregulated transcripts encoding protective proteins (desiccation-related proteins, LEAs, defensins), several matrix-modifying enzymes (β-glucosidases and β-xylosidases-encoding transcripts), specific structural proteins (AGPs; HpRGPs; Leucine-Rich Repeat Extensins – LRXs) and proteases (subtilisin-like, aspartic proteinases) **(Supplementary Table S2).**

Initial rehydration (R1 relative to DL) triggered a rapid shift towards CW restructuring and protein turnover (**Figure 3, Supplementary Table S2**). Majority of upregulated DEGs were transcripts encoding pectin-degrading and modifying enzymes (including six PGs; four PLs; four PMEIs; one PME; one pectin acetylesterase, PAE; and a PG-inhibiting protein, PGIP) and CAZymes (β-galactosidases, β-D-xylosidases, XTHs), indicating active CW relaxation. Notably, a surge in subtilisin-like proteases (13 transcripts), PODs, expansins, and AGPs was consistent with increased matrix-remodelling. Conversely, transcripts suppressed at R1 vs. DL included phenylpropanoid/lignin-related enzymes (caffeoylshikimate esterase, MODIFYING WALL LIGNIN), desiccation-protective proteins, and stress-induced CAZymes.

By R2, transcriptomic activity attenuated, limited to suppressing stress signals (dirigent proteins, PODs) and fine-tuning glycans via upregulated β-xylosidases and XTHs. No CW-associated DEGs were detected in R2 vs. R3. Across the six functional CW-related groups (**Figure 3**), the comparison between R3 and HL revealed no upregulated DEGs in R3, with only four transcripts remaining slightly elevated in HL (including PMEIs and a LEA protein). This indicates that the transcriptomic profile largely returned to baseline within 48 h (**Supplementary Table S2**).

### 3.4. Quantitative CW Proteomics Across Desiccation and Rehydration

A total of 9,399 peptide-*m/z* features were identified, corresponding to 2,812 proteins. Following selective targeting of apoplastic components, removal of non-CW proteins, and pre-normalisation data filtering and processing, 1077 peptide-*m/z* features and 354 proteins remained **(Supplementary Table 3)**.

To reduce between-run intensity differences across all 28 injections, log10-transformed abundance profiles were normalised via median alignment. Post-normalisation violin plots confirmed successful harmonisation across all six experimental conditions (bulk, HL, DL, R1, R2, R3), converging to a uniform median log10 intensity of ∼3.9 (**Supplementary Figure S3**).

In the individual 5-condition PCA model (**Figure 4a**), the first two axes accounted for 44.3% of total variance (Axis 1: 28.9%, Axis 2: 15.4%). Sample distribution was characterised by diagonal trajectories rather than non-overlapping clusters, with clear diagonal vectors separating HL vs. R1 and R1 vs. R2.

**Figure 4.**
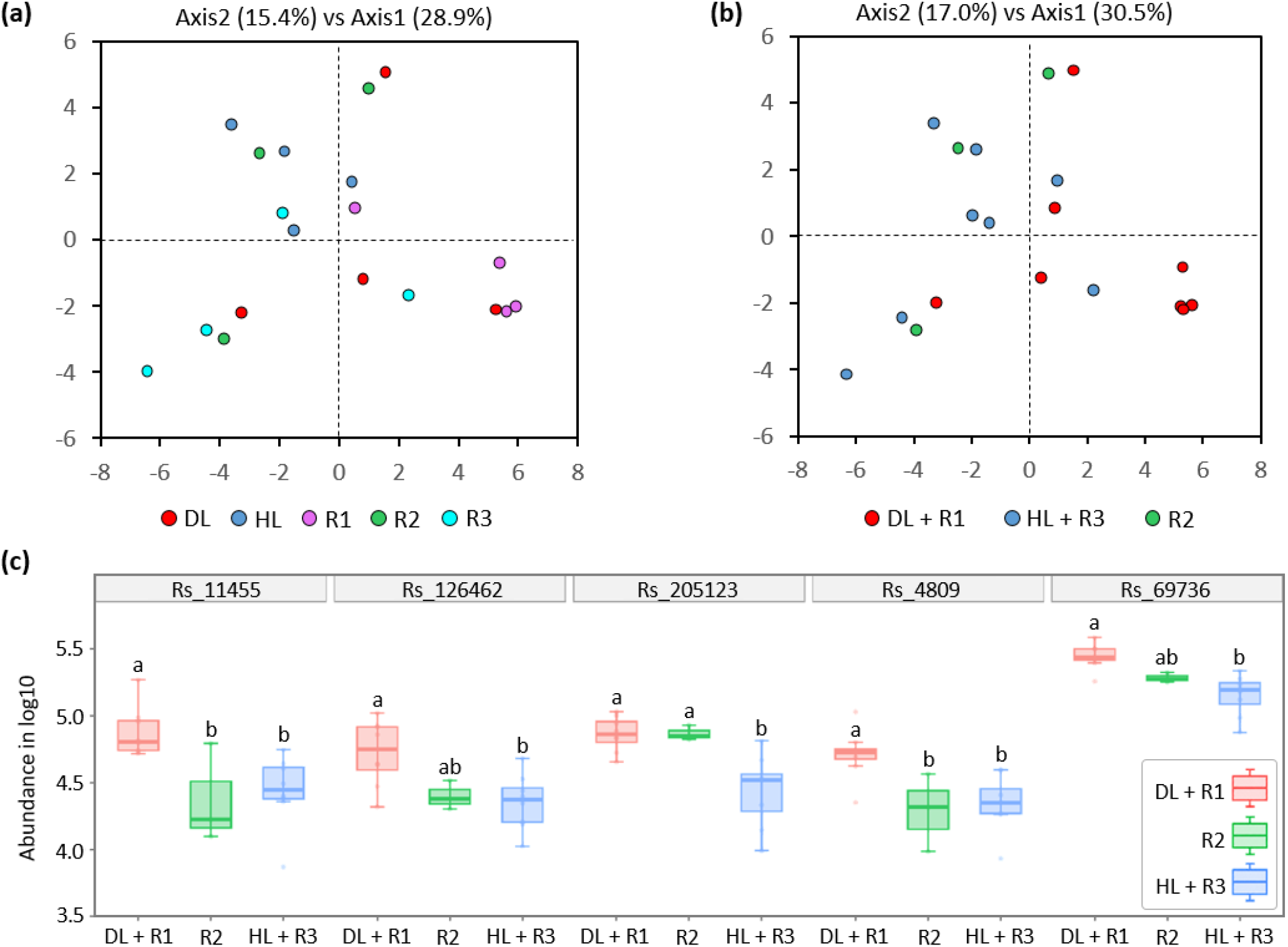
CW proteome dynamics across the desiccation-rehydration cycle. **(a)** PCA score plot of five individual states. **(b)** PCA score plot of merged tiers (DL+R1, R2, HL+R3) **(c)** Relative abundance profiles (mean ± SD) of five statistically significant DAPs across merged tiers: basic 7S globulin (Rs_11455), α-galactosidase 1 (Rs_126462), LEA4 family member (Rs_205123), miraculin (Rs_4809) and LEA4 family member (Rs_69736). Different letters indicate significant differences among merged tiers (*p*_adj_ < 0.05, one-way ANOVA with post-hoc Tukey’s HSD test).

To mitigate replicate-level noise and evaluate broader physiological patterns, conditions were merged into three functional tiers: DL+R1, R2, and HL+R3 (**Figure 4b**; 47.5% total variance; Axis 1: 30.5%, Axis 2: 17.0%). The merged model displayed a continuous trend along the diagonal axis: the DL+R1 stress tier shifted towards the lower-right quadrant, the fully recovered baseline (HL+R3) positioned in the upper-left. These multivariate patterns indicated overlapping CW proteome profiles across physiological states rather than sharp discrete state shifts during the desiccation-rehydration cycle. In addition, the hierarchical clustering reflected substantial biological variability across individual replicates (**Supplementary Figures S4,S5**). The distribution of protein coefficients of variation peaked between 35% and 45% (**Supplementary Figure S3**). Among the most highly variable proteins (CV > 70%), 7 were found at DL+R1, 4 at R2, and 4 at HL+R3.

In total, after post-normalisation data filtering and processing, 148 peptide-*m/z* features and 43 proteins remained **(Supplementary Table 3)**.

Post-hoc Tukey tests on merged tiers (DL+R1, R2, HL+R3) identified 5 key CW DAPs (*p*_adj_ < 0.05; **Figure 4c; Supplementary Table S4**): basic 7S globulin (B7SP, an extracellular xylanase inhibitor-like glycoprotein; Rs_11455), α-galactosidase 1 (Rs_126462), two LEA4 proteins (Rs_205123, Rs_69736), and miraculin (a Kunitz-type protease inhibitor; Rs_4809). Except for Rs_205123, all DAPs mapped to the same cluster (Cluster 2, **Supplementary Figure S5)**, exhibiting sharp, transient accumulation during active stress (DL+R1) followed by a rapid decline to baseline in R2 and HL+R3. Both LEA4 proteins displayed marked stress-induced accumulation and highly predicted intrinsic disorder with a strong propensity to fold into α-helices (Supplementary Figure S6). Rs_205123 maintained elevated levels across DL+R1 and R2 compared to HL+R3 (padj = 0.016), whereas Rs_69736 showed a stepwise decline from its peak in DL+R1 (*p*_adj_ = 0.016) down to baseline in HL+R3.

### 3.5. Changes in CW Polysaccharide Composition and Conformational Transitions

FTIR spectra of isolated CW material were collected in the mid-infrared region (4000–400 cm⁻¹) (**Supplementary Figure S7**). Further analysis focused on PCA of the second derivative of the polysaccharide fingerprint region (1185–870 cm⁻¹) revealing IR vibrations responsible for separation of investigated physiological states (**Figure 5a**).

**Figure 5.**
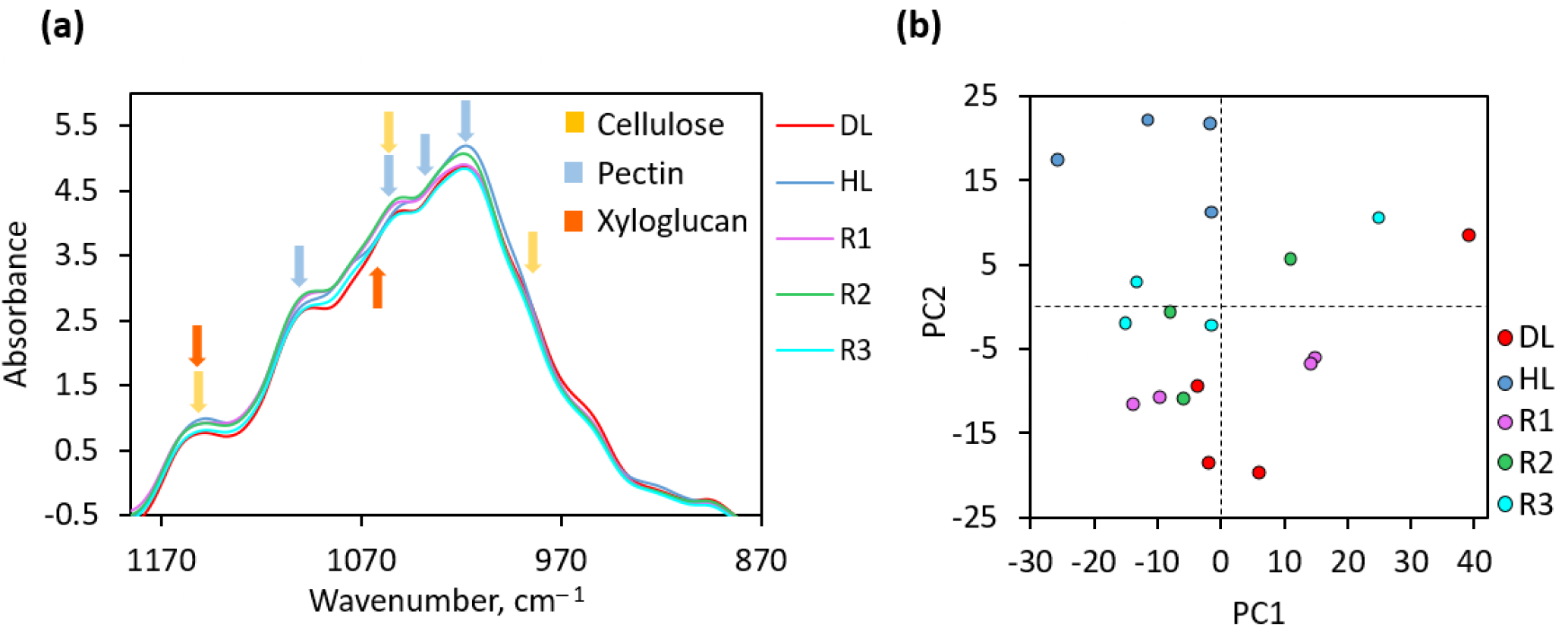
FTIR spectroscopic characterisation of purified CWs across five desiccation/rehydration stages. **(a)** Averaged, baseline-corrected, and normalised FTIR spectra of the selected statistically significant region (1185–870 cm⁻¹); arrows indicate characteristic vibrational bands assigned to cellulose (yellow), pectin (blue), and xyloglucan (orange). **(b)** PCA score plot illustrating structural trajectories and spatial separation of CW carbohydrates along PC1 and PC2, showing distinct clustering between HL, DL, R1, R2, R3.

This spectral region encompasses characteristic vibrational modes originating from main cell wall structural polymers, including cellulose (1155, 1050, 1020, and 990 cm⁻¹), xyloglucans (1150 and 1123 cm⁻¹), and various pectic domains (1092, 1065, 1036, and 1010 cm⁻¹) (**Supplementary Table S6**). Unsupervised PCA conducted on the pre-processed FTIR spectra (second derivative) clearly discriminated physiological phases along PC2 into three distinct clusters: HL (positive PC2), DL+R1 (negative PC2), and R2 with R3 (intermediate PC2 near the origin), while PC1 captured mostly HL vs DL differences (**Figure 5b, Supplementary Figure S7**). Analysis of PC2 loadings identified specific vibrational frequencies driving separation along the desiccation–rehydration trajectory. Positive PC2 loadings characterised HL, negative loadings were enriched in DL+R1, and late recovery stages (R2, R3) exhibited transitional spectral profiles returning towards baseline.

Discriminative spectral markers for the HL group (positive PC2 loadings) were identified at 1135, 1092, 1065, 1036, 1010, and 893 cm⁻¹ **(Supplementary Table S6, Supplementary Figure S7**). The prominent bands at 1092, 1065, and 1036 cm⁻¹ correspond primarily to skeletal C–O and C–C stretching and out-of-plane bending vibrations of the pectic polysaccharide backbone (Kačuráková and Wilson, 2001; Vidović et al., 2022). The band at 893 cm⁻¹ is assigned to β-glycosidic linkages in hemicellulose with contributions from amorphous cellulose fibres. Conversely, key spectral markers characterising the desiccated DL and R1 states (negative PC2 loadings) shifted to 1153, 1123, 1106, 1080, 1050, 1020, 992, and 905 cm⁻¹ (**Supplementary Figure S7, Supplementary Table S6**). The bands at 1153, 1106, 1050, and 1020 cm⁻¹ represent C–O–C asymmetric stretching and C–OH secondary alcohol vibrations of crystalline and semi-crystalline cellulose microfibrils (Vidović et al., 2022; Milić et al., 2023). Notably, the distinct band at 1123 cm⁻¹ represents a well-established marker for xyloglucan side-chains and polymer-polymer hydrogen bonding.

Overall, FTIR spectral shifts indicated reversible alterations in CW polysaccharide organisation during desiccation—characterised by a transient suppression of hydrated pectic modes—which progressively returned to control-like levels by 48 h (R3).

### 3.6. Cell-Wall-Bound Phenolic Compound Dynamics Across Desiccation and Rehydration

HPLC-DAD analysis of cell-wall-bound phenolics released via alkaline hydrolysis identified *p*-CA and FA as predominant hydroxycinnamic acids (HCA, **Figure 6**). Desiccation induced a marked accumulation of CW-bound phenolics, with total *p*-CA and FA levels increasing more than twice in DL and R1 relative to HL control levels. As recovery progressed through R2 and R3, phenolic concentrations rapidly declined compared to DL and R1, returning almost to baseline levels (no significant differences, **Figure 6**).

**Figure 6.**
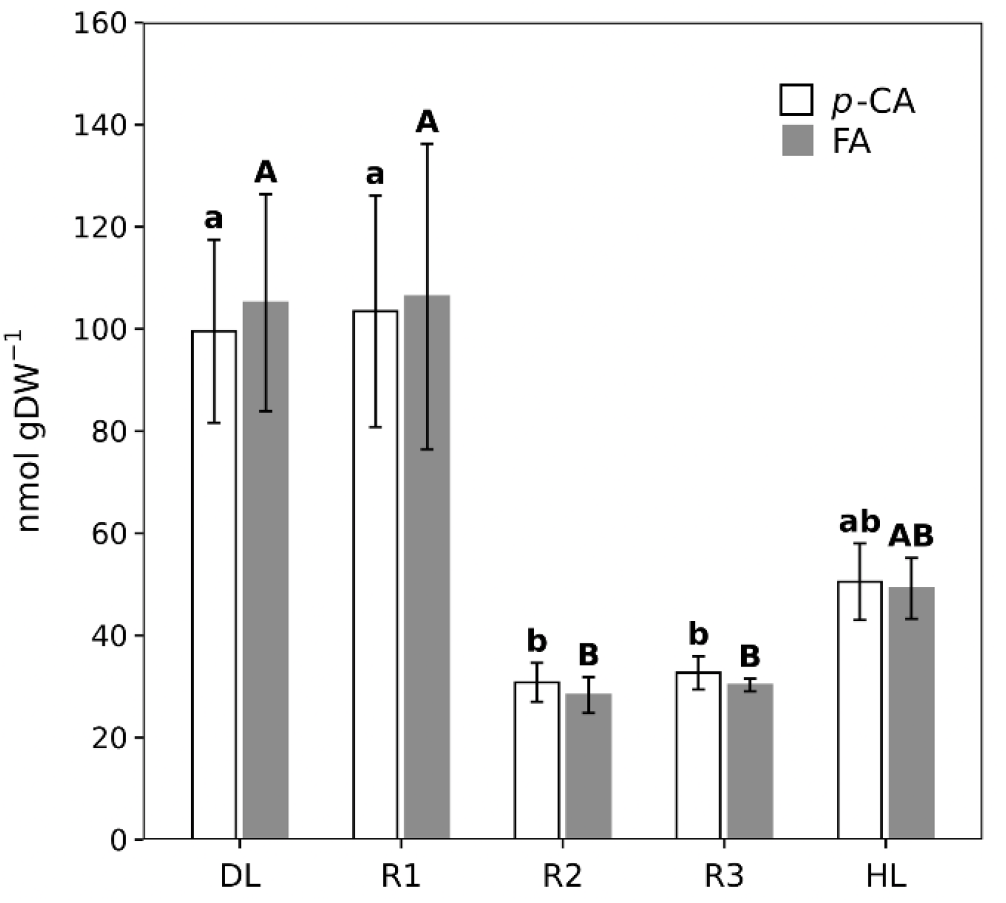
Dynamics of CW-bound *p*-coumaric acid (*p*-CA, white bars) and ferulic acid (FA, grey bars) contents in leaves across five desiccation/rehydration stages. Values are expressed as mean ± standard error (SE) in nmol gDW^−1^ (n = 4). Different lowercase (a–b) and uppercase (A–B) letters denote statistically significant differences among stages for *p*-CA and FA, respectively (*p* < 0.05, one-way ANOVA with Tukey’s HSD test).

## 4. Discussion

Vegetative desiccation tolerance relies on biomechanical strategies that prevent irreversible CW disrupting and preserve plasma membrane integrity as turgor decline. Expanding beyond the preliminary molecular snapshot of desiccation-induced CW changes *in R. serbica* (Vidović et al. 2022), here we provide novel insights into the spatiotemporal structural reorganisation of the CW components that maintain their integrity during dehydration and recovery.

### 4.1. Desiccation

FTIR profiling confirmed a decrease in pectin, cellulose, hemicellulose, and xyloglucan levels during desiccation, consistent with our previous observations (Vidović et al. 2022). Unlike *C. plantagineum*, where ELISA showed an overall reduction in LM20 binding and increase in LM19 (Jung et al. 2019), *R. serbica* retains both high-methylated (HG-HM; LM20) and low-methylated (HG-LM; LM19) homogalacturonan pools, with spatially differentiated labelling patterns (microdomains) that may contribute to local mechanical heterogeneity. HG-HM at convolutions might provide matrix flexibility to accommodate deep folding without structural disruption. Conversely, HG-LM can form a regulatory node: blockwise demethylation by PMEs promotes Ca^2+^-crosslinking to reinforce wall stability under high stress, whereas random demethylation targets HGs for cleavage by endo-polygalacturonases and pectate lyases, facilitating localised wall loosening (Hocq et al. 2017; Wormit & Usadel, 2018). Simultaneously, increased detection of LM16-recognised RG-I-associated epitopes can act as hydration-retaining plasticisers that prevent irreversible cellulose microfibril aggregation (Harholt et al. 2010). The detection of PME transcripts alongside widespread suppression of CW-loosening CAZymes, expansins, and GRPs/PRPs upon full desiccation indicates that this dual pectin modulation was likely established earlier, during early-to-moderate dehydration.

In desiccated *R. serbica* leaves, labelling for extensins (LM1) and HpRGPs—specifically AGPs-associated, LM2 labelling exhibited distinct vesicle-like structures—extended throughout the cell interior. A similar structural strategy was observed in dicotyledonous resurrection plants (*C. plantagineum*, *M. flabellifolia*), where dynamic alterations in xyloglucan composition and pectin methylation regulate matrix plasticity and enable controlled CW folding without mechanical injury (Vicré et al. 2004; Moore et al. 2008; Jung et al. 2019). Many transcripts encoding structural CW proteins were downregulated in DL (e.g., expansins, GRPs, PRPs, and HpRGPs), although selected members (AGPs; HpRGPs; LRXs) showed the opposite response. Nevertheless, pre-existing structural protein networks—as documented for CpGRP1 in *C. plantagineum* and BhGRP1 in *Boea hygrometrica* (Wang et al. 2009; Giarola et al. 2016)—maintain CW integrity. Parallel to these physical compactions, desiccation induces a tight spatial condensation of the structural cellulose-hemicellulose framework and an accumulation of CW-bound *p*-CA and FA. The increased CW-bound phenolic content is consistent with a potential contribution to wall reinforcement, although the specific polymeric attachment sites and the extent of oxidative cross-linking were not directly determined (Mnich et al. 2020).

Targeted induction of desiccation-related and defensin-like genes, emphasises the importance of establishing an apoplastic barrier against mechanical rupture and pathogens, similar to mechanisms described in other resurrection plants (Gechev et al. 2012; Shumayla et al. 2025). Moreover, two DAPs elevated in DL+R1 belonged to the LEA4 protein family. Although LEA proteins are typically considered intracellular molecular shields (Tunnacliffe & Wise, 2007), this is the first report confirming their presence in the ionically bound CW fraction of a resurrection plant. Based on *in vitro* results for Rs_LEA30 (Pantelić et al. 2025), we hypothesise that these two LEA4 protein family members transition from intrinsically disordered states in hydrated environment to amphipathic α-helices upon desiccation, where they can protect apoplastic enzymes from denaturation by masking hydrophobic or charged patches, while their positively charged faces might electrostatically interact with COO^−^ groups of demethylesterified pectins to stabilise CW matrix during DL and R1. Additionally, LEA proteins may form protective hydrogels or biomolecular condensates and contribute to reactive oxygen species (ROS) scavenging or protective molecular shielding (Hundertmark & Hincha, 2008; Olvera-Carrillo et al. 2010; Graether & Boddington, 2014; Belott et al. 2020).

### 4.2. Short-term response to rewatering

Rehydration represents a highly vulnerable transitional phase susceptible to cellular oxidative damage before intracellular antioxidative systems fully recover (Sgherri et al. 1994). To counteract this threat, the sustained abundance of desiccation-induced CW DAPs through 1 h post-rewatering (R1) may help preserve apoplastic protein protection and structural integrity. Additionally, CW-bound HCAs remain elevated at R1, potentially contributing dual biomechanical and antioxidant roles by regulating apoplastic mechanical strength and ROS dynamics (Mnich et al. 2020). Transcriptomic profiling of R1 revealed the specific induction of two POD isoforms (sharing 97% sequence identity) annotated as *Arabidopsis thaliana* Peroxidases 42 (AtPrx42; **Supplementary Table S2**). These cationic isoforms (calculated pI 8.52) align with previously reported cationic CW-bound PODs (pI 8.4–9.3) induced during *R. serbica* rehydration (Veljović-Jovanović et al. 2006). These ionically bound CW enzymes exhibit a high substrate affinity for HCAs such as FA (Km = 1.9 mM), suggesting that they scavenge apoplastic H_2_O_2_ and polymerise FA to reinforce the CW. The apoplastic redox state at R1 is further influenced by the rapid suppression of ascorbate oxidase (AO), which directly controls reduced ascorbate availability (Mellidou & Kanellis, 2024). Combined with the cationic PODs induction, this AO downregulation could favour the maintenance of reduced ascorbate levels (reported in Sgherri et al. 2004), which can directly scavenge H_2_O_2_ and recycle oxidised HCA derivatives and contribute to POD-mediated H₂O₂ removal through the reduction of oxidised phenolic intermediates (Takahama, 2004; Veljović-Jovanović et al. 2006). By regulating H_2_O_2_ levels, the generation of highly reactive hydroxyl radicals, HO^•^ (e.g., via Fenton/Haber–Weiss reactions) is controlled, thereby potentially affecting HO^•^–mediated CW polysaccharides cleavage and site-specific CW loosening and expansion (Schopfer, 2001; Fry, 2004).

In parallel with this protective and redox-controlled backdrop, *R. serbica* leaves at R1 rapidly activate a CW remodelling programme to support swift cell expansion. A sudden decrease in HG-HM (LM20) labelling coincides with a continuous HG-LM (LM19) lining along expanding cell borders and the upregulation of genes encoding pectin-modifying enzymes at R1 relative to DL. While upregulated *PMEs* can promote controlled demethylesterification to reinforce CW rigidity against osmotic swelling, co-induced *PMEIs, PLs, PGs* and their inhibitors (*PGIPs*) may locally regulate pectin cleavage and Ca^2+^-dependent stiffening during CW expansion needed for rapid turgor recovery. The increased abundance of α-galactosidase (Rs_126462) in DL+R1 may contribute to CW-associated carbohydrate turnover, although its endogenous substrates in *R. serbica* remain unidentified. It is known that apoplastic α-galactosidase can cleave terminal α-galactosyl residues from pectin side-chains (such as RG-II side chain A and terminal caps of RG-I/AGPs), loosening the pectic network and exposing polysaccharide substrates for downstream remodelling enzymes (Chrost et al. 2007; Moore et al., 2008). These rapid pectic events extend the model proposed for *C. plantagineum*, where de-esterified HGs act as docking platforms for apoplastic CpGRP1 and wall-associated receptor kinases (CpWAK1)—to coordinate stress sensing and CW integrity (Jung et al. 2019; Chen et al. 2021). Notably, the differential expression of specific WAKs between DL and R1 in *R. serbica* **(Supplementary Table S2)** suggests a possible link between pectin remodelling and CW-associated signalling during rehydration.

Although pectin-modifying enzymes were induced at the transcript level, they were not identified as DAPs in the CW proteome. Given that these enzymes were detected in the raw proteomic dataset without reaching statistical significance, this transcript–protein discrepancy may reflect a delay between transcript induction and protein accumulation, localised changes in enzymatic activity, or technical limitations in proteomic coverage.

Besides pectins, at LM2-detected AGP-associated punctate signals, transiently reorganises along expanding cell boundaries at R1. This altered spatial distribution, reflects targeted exocytosis and endosomal recycling during localised CW assembly (Nguema-Ona et al. 2012; Lampugnani et al. 2024). The co-induction of *LRXs* and structural glycoproteins at R1 may contribute to the reorganisation of CW-associated protein networks during expansion. In parallel, early-rewatering-induced genes encoding subtilases facilitate apoplastic signal transduction, protein turnover, and the proteolytic activation of precursor enzymes and CW-modifying proteins (Schaller et al. 2018). Together with CAZyme upregulation, this transcriptional response suggests increased capacity for apoplastic protein turnover to enable controlled CW expansion upon turgor recovery.

### 4.3. Recovery

Upon *R. serbica* rehydration (R2–R3), the CW matrix progressively relaxes and recovers, moving back to functional homeostasis, as suggested by the intermediate positioning of R2 and R3 samples along PC2 and the accompanying changes in pectin-associated spectral features. Concurrently, CW-bound *p*-CA and FA levels dropped almost threefold between R1 and R2, declining to levels slightly below HL controls by R2– R3. This reduction is consistent with the general downregulation of phenylpropanoid and lignin biosynthetic genes during rehydration.

This reduction in extractable CW-bound phenolics accompanying CW structural recovery occurs simultaneously with dynamic pectic and glycoprotein reorganisation. Across R2 and R3, the restoration of LM20 and LM19 signals, alongside the spatial relaxation of RG-I galactans (LM16), is consistent with rehydration and reorganisation of the pectic matrix. As in *C. plantagineum* and *M. flabellifolia*, restoring CW flexibility relies on replacing rigid, Ca^2+^-chelated HG-LM with newly synthesised HG-HM, while RG-I galactan side-chains function as molecular plasticisers that maintain matrix fluidity (Vicré et al. 2004; Moore et al. 2006, 2013; Jung et al. 2019). Simultaneously, structural CW glycoproteins re-establish their homeostatic distribution. Functioning potentially as biomechanical buffers during osmotic transitions (Seifert & Roberts, 2007), AGP signals that began reorganising at R1 continue to redistribute through R2 and R3 into continuous lines along the plasma membrane–CW interface, as observed in HL, consistent with recovery of this cellular boundary. Likewise, the condensed extensin network relaxes into a regular, three-dimensional scaffold, providing the structural flexibility needed to sustain fully turgid cells.

Remarkably, transcriptomic profiles diverge minimally between R2 and R3, with virtually no CW-targeted DEGs detected in R3 compared to fully HL. This attenuated molecular activity indicates that the biggest part of the observed enzymatic and structural remodelling occurs within the first day of rewatering (R1– R2). Together, these findings support substantial structural and molecular recovery of the CW in *R. serbica* within 48 h of rewatering, aligning with other resurrection angiosperms like *C. plantagineum* and *B. hygrometrica* (Vicré et al., 2004; Zhu et al., 2015).

## 5. Conclusion

Based on our multi-omics, spectroscopic, and immunocytochemical findings, we propose an integrated model of apoplastic biomechanical adaptation and recovery in *R. serbica* (**Figure 7**).

**Figure 7.**
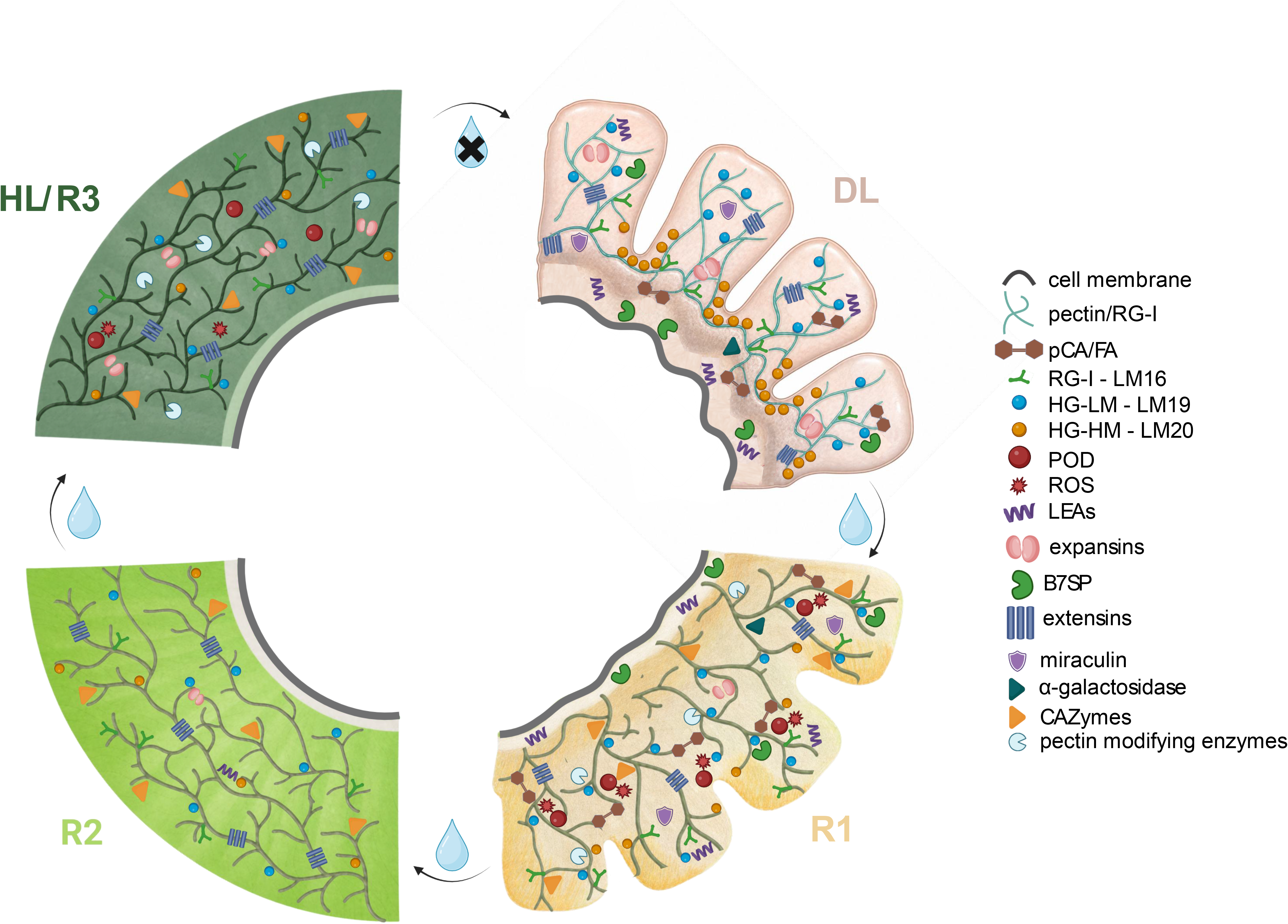
Proposed model of CW flexibility and stabilisation in *R. serbica*. Starting from hydrated homeostasis (HL), severe desiccation (DL) induces controlled CW folding supported by highly methylesterified homogalacturonans (HG-HM) and RG-I galactans at convolutions, and low methylesterified homogalacturonans (HG-LM) alongside sell boundaries, CW-bound *p*-coumaric and ferulic acid (*p*-CA/FA), and ionically bound CW LEA4 family proteins and basic 7S globulin (B7SP). Initial rewatering (R1) triggers a rapid redistribution of pre-existing AGP vesicular structures along expanding cell borders (omitted from this scheme for visual clarity), alongside upregulation of transcripts encoding peroxidases (POD), and CW-remodelling enzymes (carbohydrate-active enzymes – CAZymes, proteases, and pectin modifying enzymes) that initiate matrix relaxation. During recovery (R2–R3), CW-bound phenolic levels decrease, allowing the pectic–glycoprotein matrix to fully re-hydrate and restore its baseline CW architecture within 48 h.

In fully hydrated tissues (HL), the CW exhibits a relaxed architecture. Upon desiccation, controlled CW folding is associated with spatial differences in HG methylesterification: highly methylesterified HGs and RG-I galactans preserve matrix fluidity at convolutions to enable deep tissue folding, whereas demethylesterified HGs line cell borders to provide structural rigidity where mechanical support is required. The spatial organisation of AGP epitopes and condensation of extensin-associated structures may help preserve the plasma membrane–CW interface, while increased CW-bound *p*-CA and FA levels may contribute to matrix reinforcement. Simultaneously, LEA4 proteins accumulate in the CW fraction and may protect CW-associated biomolecules during desiccation, alongside B7SP and miraculin, which may provide additional protection against enzymatic CW degradation (**Figure 7**).

Upon acute rewatering (R1), initial water influx triggers a rapid transition characterised by the redistribution of AGP-associated vesicular structures towards expanding cell borders. This is accompanied by increased expression of transcripts encoding CW-remodelling enzymes (PMEs, PGs, PLs, CAZymes, proteases) potentially contributing to localised matrix relaxation, alongside ROS scavenging and phenolic cross-linking control via PODs. As rehydration advances through structural restoration (R2) to restored homeostasis (R3), CW-bound *p*-CA and FA contents decrease, alongside the recovery of pectic– glycoprotein hydrogel and extensin scaffolds that re-expand into their baseline architecture. Together, these findings support substantial structural and molecular recovery of the CW within 48 h of rewatering (**Figure 7**).

## Supporting information

SUPPLEMENTARY FIGURES Supplementary Figure S1. Tissue-specific cellular responses and cross-sectional area dynamics across desiccation and rehydration

## Supporting Information

Additional supporting information can be found online in the Supporting Information section.

**Supplementary Figure S1.** Tissue-specific cellular responses and cross-sectional area dynamics across desiccation and rehydration stages in *Ramonda serbica*. Data in the box plot represent median values (horizontal lines), mean values (open triangles), upper and lower quartiles (boxes), and individual cell measurements (n ≥ 30 cells per condition; grey dots). Percentages indicate mean area relative to the initial HL state (100%). Scale bars = 50 μm.

**Supplementary Figure S2.** Expression profiles of curated cell wall-targeted transcripts across hydrated control (HL), desiccated leaves (DL), initial rehydration (R1, 1 h), mid-rehydration (R2, 24 h), and fully rehydrated leaves (R3, 48 h). Z-score normalised expression values are colour-coded (red: higher relative expression; blue: lower relative expression). Dendrograms on the left depict hierarchical clustering based on expression pattern similarities, highlighting distinct stage-specific transcript clusters.

**Supplementary Figure S3.** Comprehensive quality control assessment of raw extraction ion chromatogram (XIC) LC-MS/MS data prior to feature filtering using the MCQR package. **(a)** Violin plots showing intensity harmonisation before and after median-based normalisation across merged tiers (DL+R1, R2, HL+R3). **(b)** CV frequency distribution showing protein variance within merged group (DL+R1, R2, HL+R3) groups protein variance **(c)** Distribution of chromatographic peak width (in seconds) across all injections, showing high peak shape consistency with median width of 18.2 s. **(d)** Distribution of log10-transformed peptide-*m/z* intensity across individual LC-MS runs, demonstrating consistent dynamic range and signal distribution. **(e)** Global distribution of standard deviation of peptide retention times (RT in seconds) across runs, highlighting overall high run-to-run reproducibility. **(f)** Zoomed-in analysis of retention time variability (SD < 100 s) of detected peptide-*m/z* features across liquid chromatography runs.

**Supplementary Figure S4.** Unsupervised two-dimensional hierarchical clustering and expression heatmap of quantified cell wall proteins across all individual biological replicates (n = 21 MS runs). Rows represent individual cell wall proteins, and columns represent individual biological replicates. Relative abundance values are colour-coded (red indicating higher abundance, blue indicating lower abundance). The upper dendrogram highlights strong biological clustering of replicates within their corresponding physiological states.

**Supplementary Figure S5.** Partitioning of 43 quantified cell wall proteins into six distinct response clusters based on k-means clustering across merged condition tiers (DL+R1: combined desiccation and early rehydration; R2: intermediate rehydration; HL+R3: fully recovered/control state). Y-axes depict mean-centred scaled abundance patterns for each cluster profile. Protein assignments for each cluster (Cluster 1 through Cluster 6) are detailed in **Supplementary Table S4**.

**Supplementary Figure S6.** *In silico* structural characterisation of desiccation-responsive cell wall LEA4 proteins. Structural bioinformatic characterisation of cell wall-associated LEA4 proteins (**a)** Rs_69736 and **(b)** Rs_205123. Structural models were predicted using AlphaFold3 (https://alphafoldserver.com/), displaying predicted pTM scores (pTM = 0.27 and 0.25, respectively). Selected segments of amphipathic α-helices (residues 135–161 for Rs_69736 and 101–119 for Rs_205123) were visualised using helical wheel projections via HeliQuest (https://heliquest.ipmc.cnrs.fr/), highlighting the clear spatial separation of hydrophobic (yellow) and charged/polar residues. Intrinsic disorder percentages were predicted using three independent bioinformatic tools: FELLS (http://old.protein.bio.unipd.it/fells/), IUPred3 (https://iupred3.elte.hu/), and NetSurfP-3.0 (https://services.healthtech.dtu.dk/services/NetSurfP-3.0/), uniformly confirming a highly intrinsically disordered character across both proteins.

**Supplementary Figure S7.** FTIR cell wall polysaccharide spectra and Principal Component Analysis (PCA) loading profiles. **(a)** FTIR spectra of the collected mid-infrared region **(b)** Loading plots for PC1 (top) and PC2 (bottom) in the 1185–870 cm⁻¹ fingerprint region. Wavenumbers (cm⁻¹) with positive PC2 values represent key spectral features discriminating the fully hydrated state (HL; e.g., pectic domain vibrations), whereas negative PC2 values denote spectral markers characteristic of the desiccated (DL) and early rehydrated (R1) states (e.g., crystalline cellulose microfibrils and xyloglucan cross-links). All assigned vibrational frequencies and their corresponding polysaccharide contributions are detailed in **Supplementary Table S6**.

**Supplementary Table S1.** Master dataset containing transcript-level differential expression results for all ten pairwise comparisons among the five physiological stages (HL, DL, R1, R2, and R3). The table includes transcript identifiers, functional annotations, normalised expression statistics, log2 fold-change values, standard errors, test statistics, raw *p*-values, and *p*_adj_ generated by DESeq2. Differentially expressed transcripts were defined using *p*_adj_ < 0.05 and |log2FC| ≥ 2.

**Supplementary Table S2.** The dataset served as the reference source for subsequent targeted filtering and functional curation of cell wall-associated transcripts.

**Supplementary Table S3.** Quantitative breakdown of peptide-m/z features, unique peptides, and protein groups retained at each sequential quality control and filtering step during the label-free XIC pipeline. The table outlines initial identification metrics, manual curation of cell wall-localised proteins, chromatographic peak filtering, median intensity normalisation, bulk sample exclusion, and minimum unique peptide criteria. Detailed protein-level annotations for manually curated cell wall candidates are provided in Sheet 2 (*Manually curated CW proteins*).

**Supplementary Table S4.** Detailed assignments of quantified cell wall proteins (n = 43) across six k-means clusters based on relative abundance dynamics across merged condition tiers (DL+R1, HL+R3, R2). Includes protein accession identifiers (accession), unique peptide codes (protein), functional BLAST annotations (descr), scaled log10 abundance values per merged condition, and designated cluster number (cluster_number).

**Supplementary Table S5.** Master dataset containing raw and normalised quantification parameters for all individual MS runs across experimental conditions (DL+R1, HL+R3, R2). The table includes raw intensity values (q), log10-transformed values (log10q), metadata variables (condm, repm, batch), and mean condition abundances. The second worksheet (*ANOVA*) provides one-way ANOVA *p*-values and *p*_adj_ for condition-dependent abundance differences.

**Supplementary Table S6.** Assignment of characteristic FTIR vibrational bands of CW polysaccharides in *R. serbica* leaves and their contribution to PCA group separation.

**Supporting Information File S1**. Xtandem parameter for protein identification.

**Supporting Information File S2**. R script for quantitative proteomics data processing and statistics.

## Acknowledgements

This work was funded by the Ministry of Science, Technological Development and Innovation of the Republic of Serbia (Contract No. 451-03-33/2026-03/200042) and Bilateral project between the Republic of Serbia and the Republic of France, programme Hubert Curien Partnership (PHC) “Pavle Savić” (VARIEGOMICS). A.P. wishes to acknowledges the possibility of her Short-Term Scientific Missions in the Institute of Agrophysics, Polish Academy of Sciences, supervised by Agata Leszczuk and financed by COST Action (CA18210) ‘Oxygen sensing novel means for biology and technology of fruit quality’ and Labena company (https://labena.rs/). Proteomics analyses were performed on the PAPPSO facility (https://doi.org/10.15454/1.5572393176364355E12) which is supported by Paris Saclay University (https://www.universite-paris-saclay.fr/en), INRAE (http://www.inrae.fr), CNRS (http://www.cnrs.fr), AgroParisTech (https://www.agroparistech.fr/en), the Ile-de-France regional council (https://www.iledefrance.fr/education-recherche), IBiSA (https://www.ibisa.net), Saclay Plant Sciences-SPS (ANR-17-EUR-0007), Plant2Pro® (#24 CARN 0024-01), and the French proteomics infrastructure (ProFI FR2048, ANR-10-INBS-08–03).

## Data Availability Statement

The data that support the findings of this study are openly available in the Zenodo data base (10.5281/zenodo.22046118). The mass spectrometry proteomics data have been deposited to the ProteomeXchange Consortium via the PRIDE partner repository with the dataset identifier PXD083564. Supporting data are provided in the supplementary materials and the repositories listed above.

