## SUPPLEMENTARY FIGURES Supplementary Figure S1. Tissue-specific cellular responses and cross-sectional area dynamics across desiccation and rehydration for "Spatiotemporal pectin remodelling, glycoproteins, and LEA proteins maintain cell wall integrity during desiccation and rehydration in *Ramonda serbica*": Supplementary_Figures.pdf

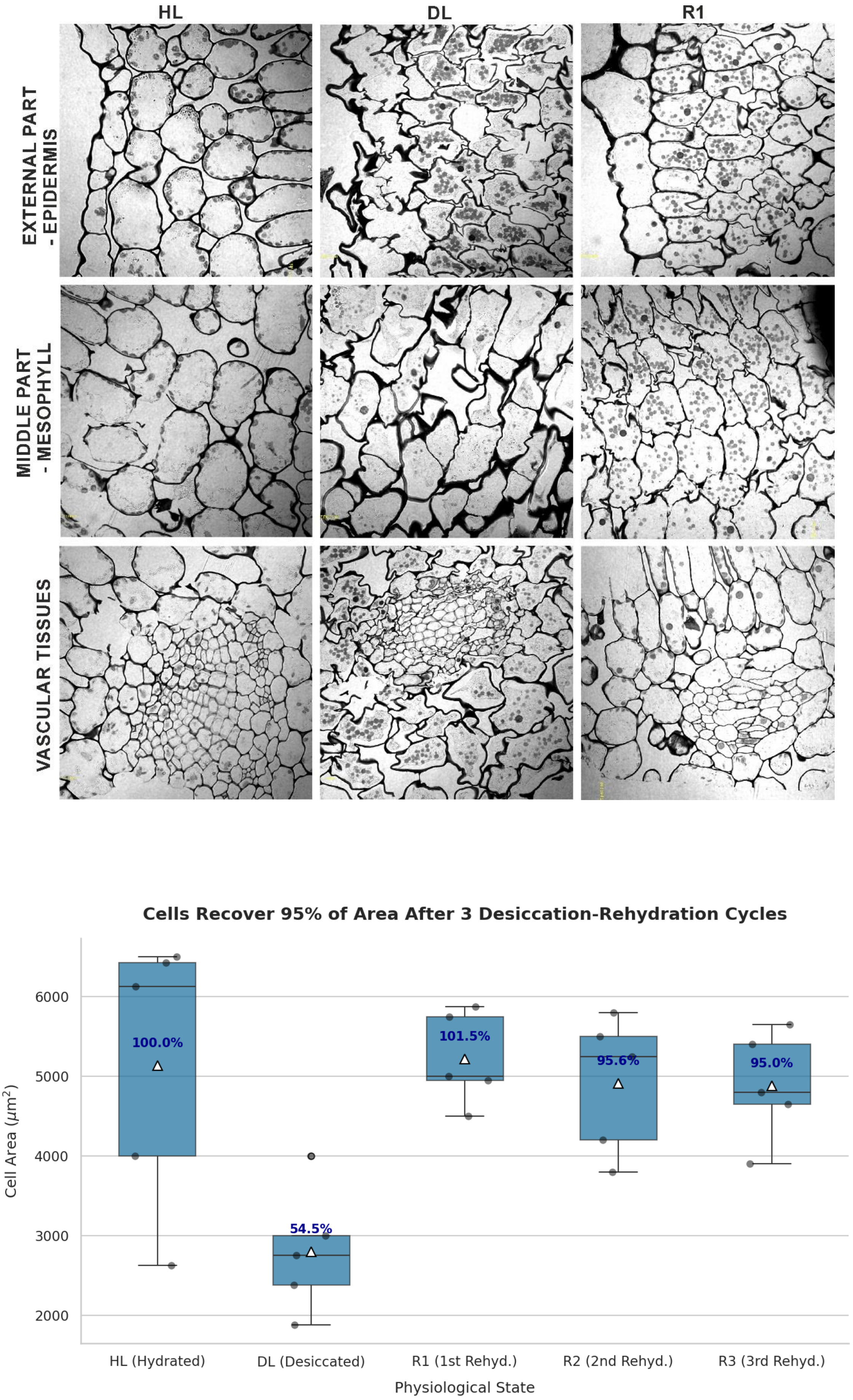


**Supplementary Figure S1.** Tissue-specific cellular responses and cross-sectional area dynamics during desiccation and rehydration cycles in *Ramonda serbica*.

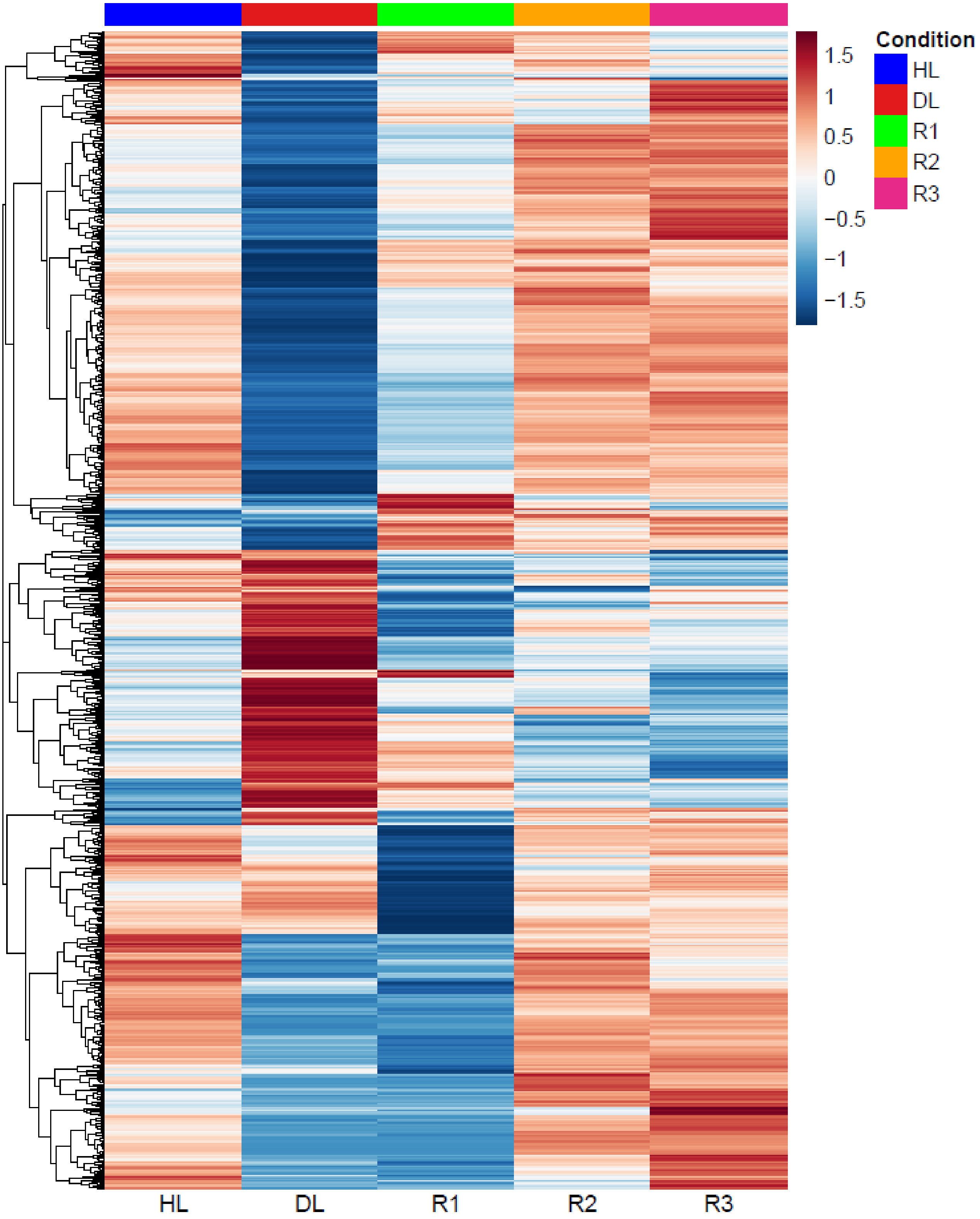


**Supplementary Figure S2.** Hierarchical clustering heatmap of cell wall-associated transcript expression across desiccation and rehydration stages in *Ramonda serbica*

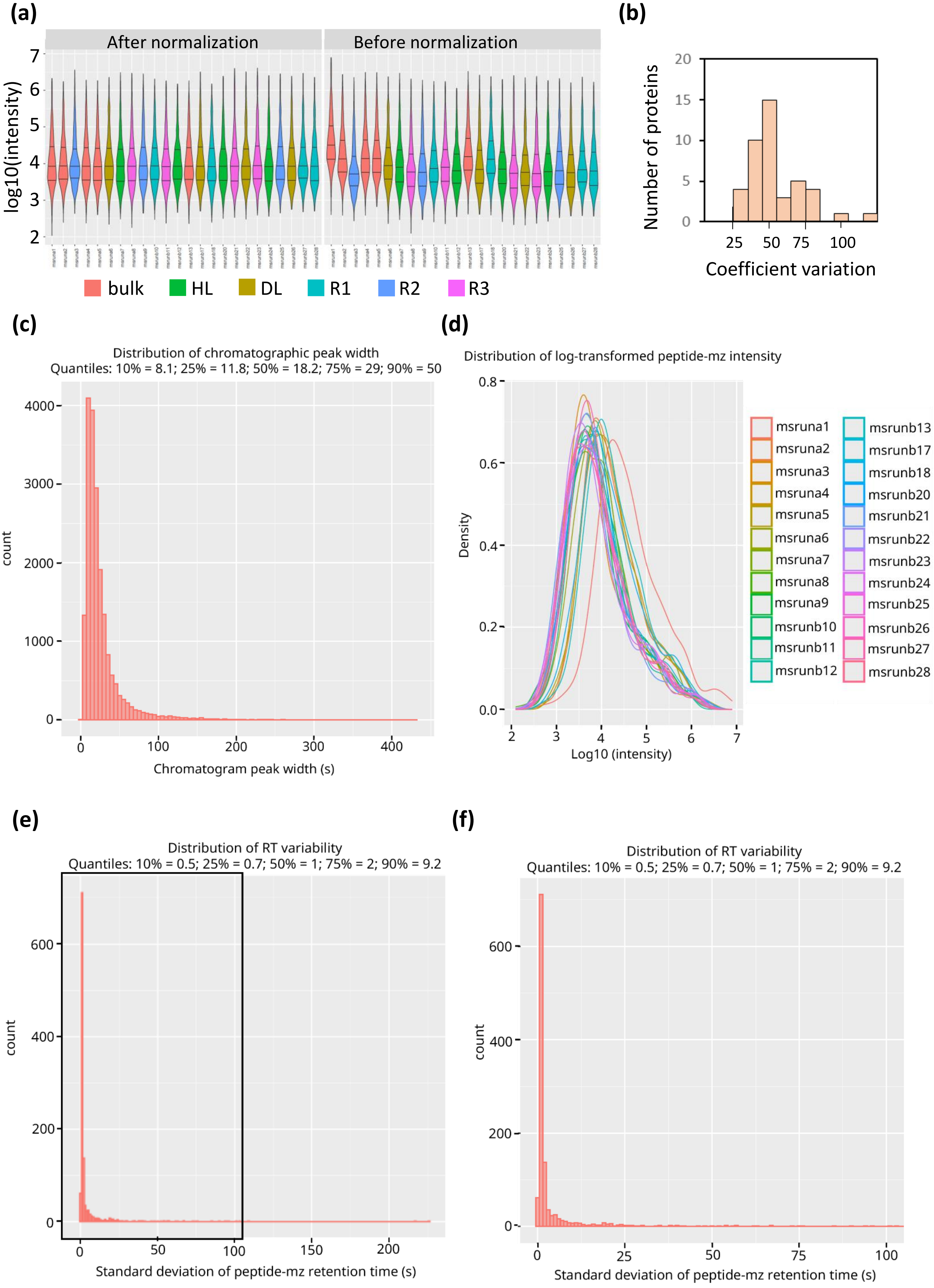


**Supplementary Figure S3.** Raw LC-MS/MS data quality control

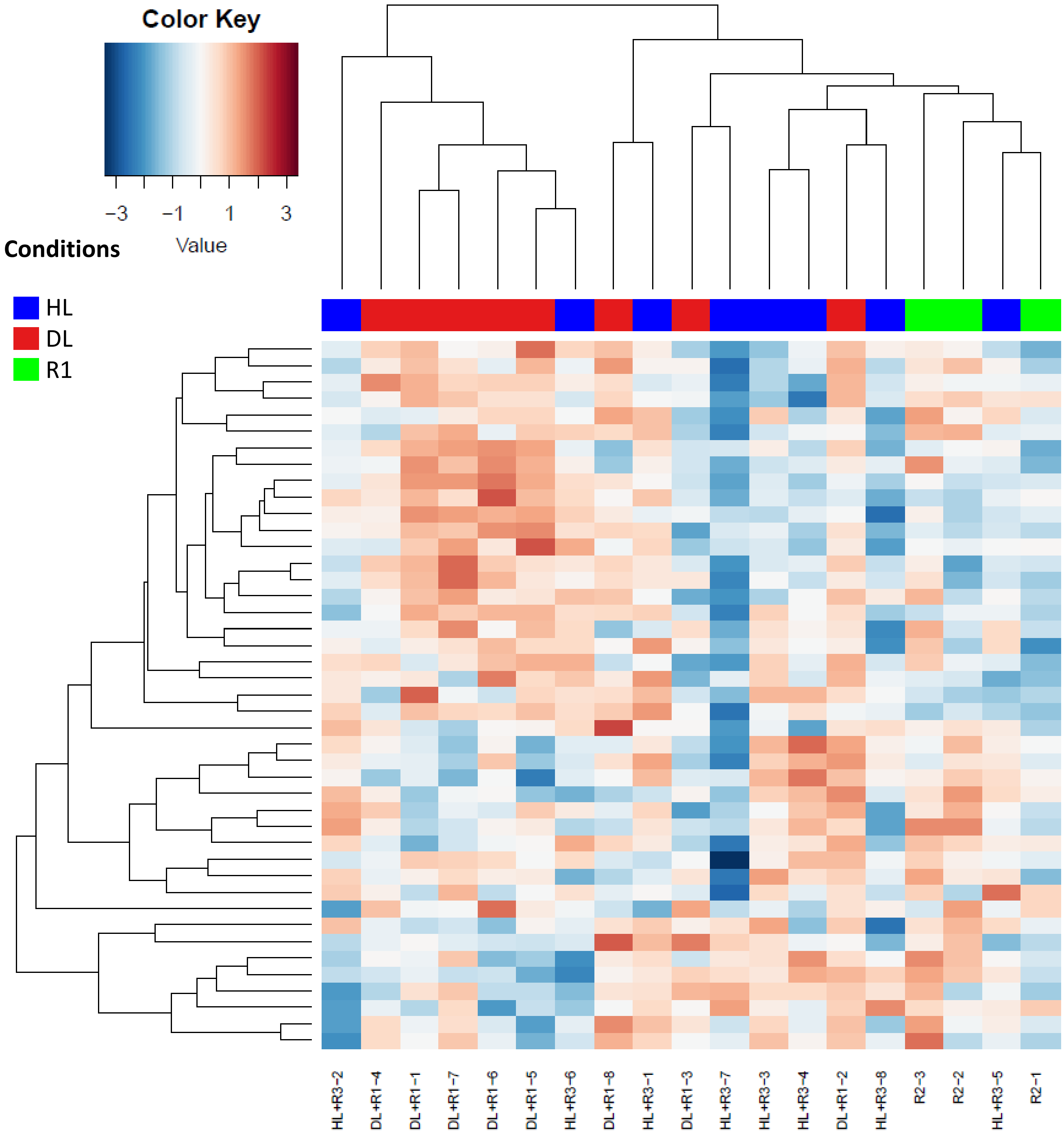


**Supplementary Figure S4.** Hierarchical clustering and expression heatmap of cell wall proteome

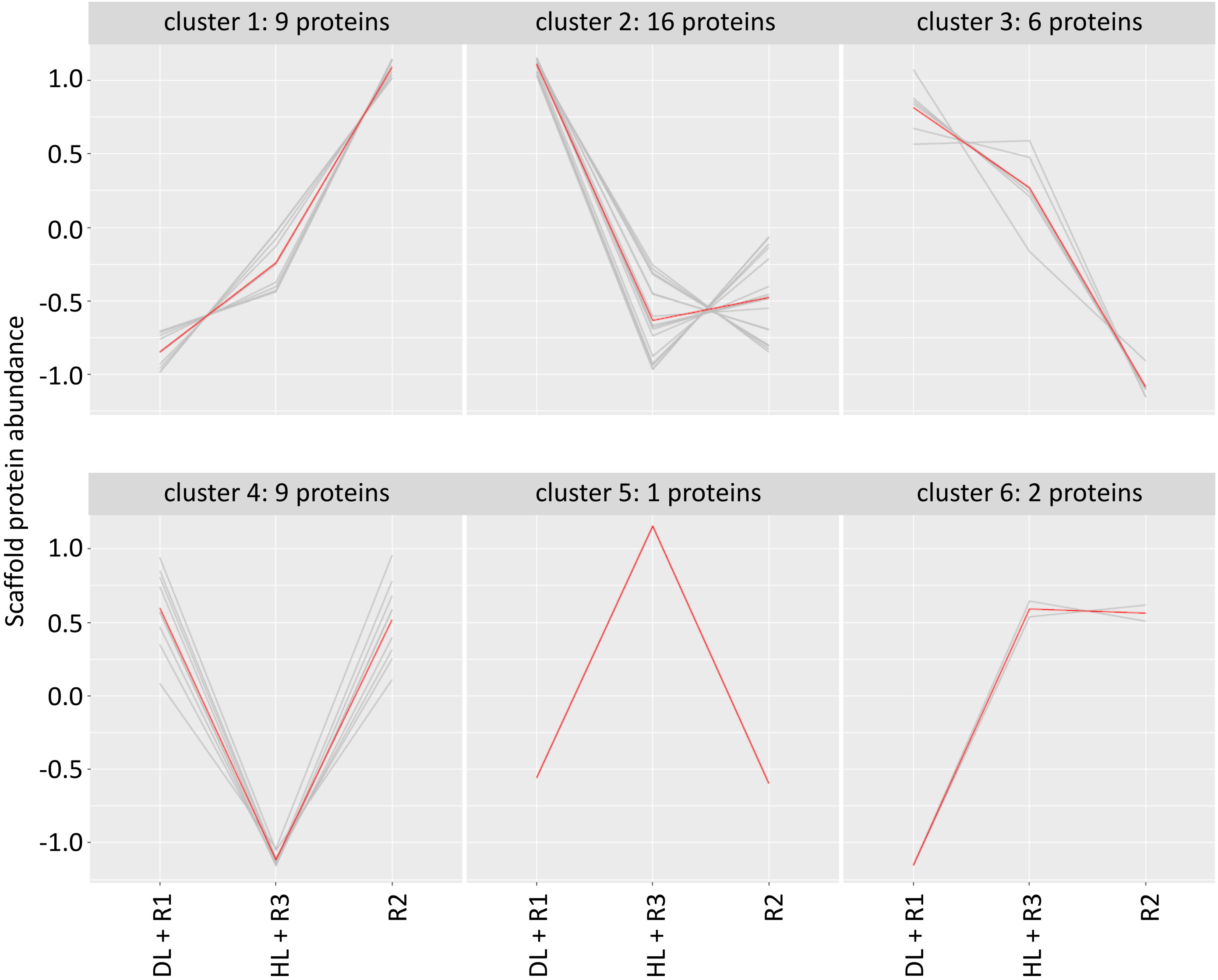


**Supplementary Figure S5.** Expression profiling and k-means clustering of the cell wall proteome across merged stress and recovery conditions.

**(a)**

**Rs_69736**

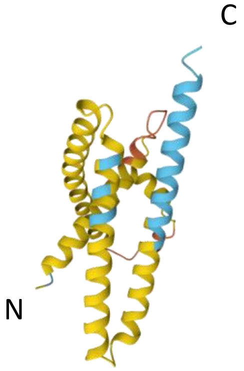


pTM = 0.27

$_{135}$DTLSWAKEKVKAGYDAAK$_{161}$

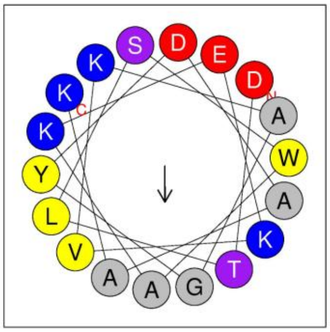

Hydrophobic face: AAVLY

| Software | Disorder (%) |
|---|---|
| FELLS | 84.86 |
| IUPred3 | 58.90 |
| NetSurfP 3.0 | 92.42 |

**(b)**

**Rs_205123**

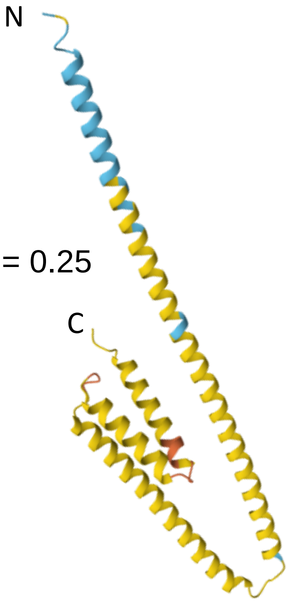


pTM = 0.25

$_{101}$VADKAKEAKDKASELGGD$_{119}$

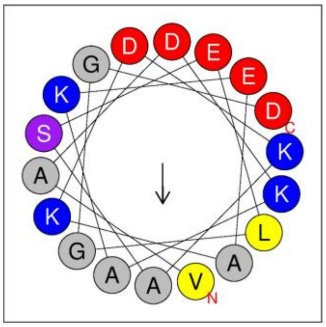

Hydrophobic face: LAFAA

| Software | Disorder (%) |
|---|---|
| FELLS | 90.77 |
| IUPred3 | 47.09 |
| NetSurfP 3.0 | 87.80 |

**Supplementary Figure S6.** *In silico* structural characterization of desiccation-responsive LEA proteins.

(a)

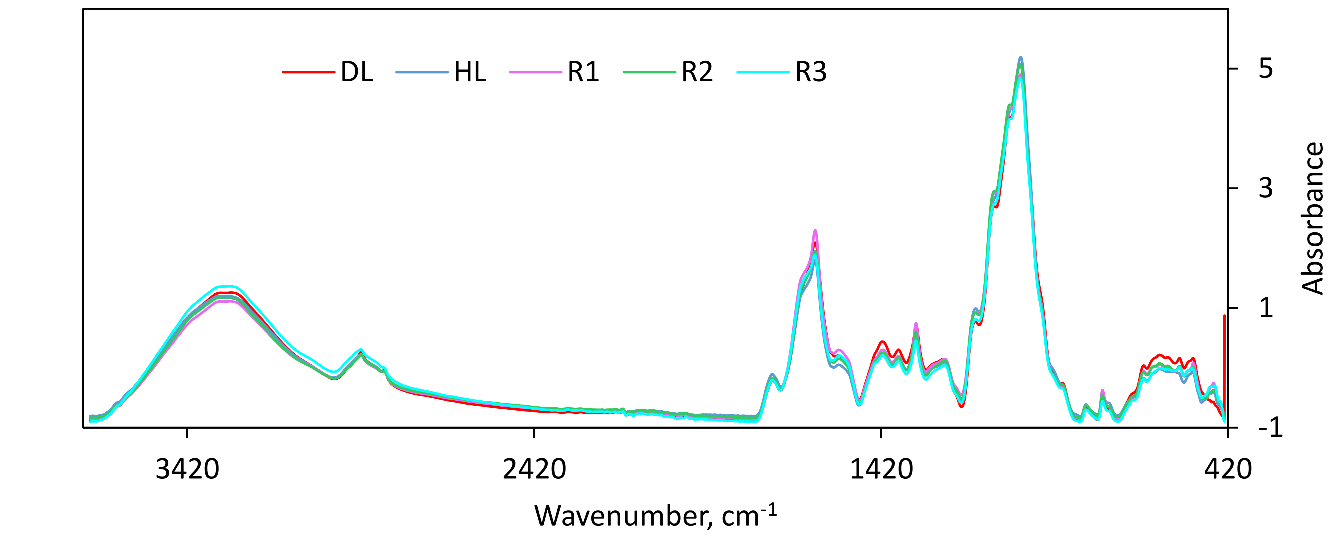


(b)

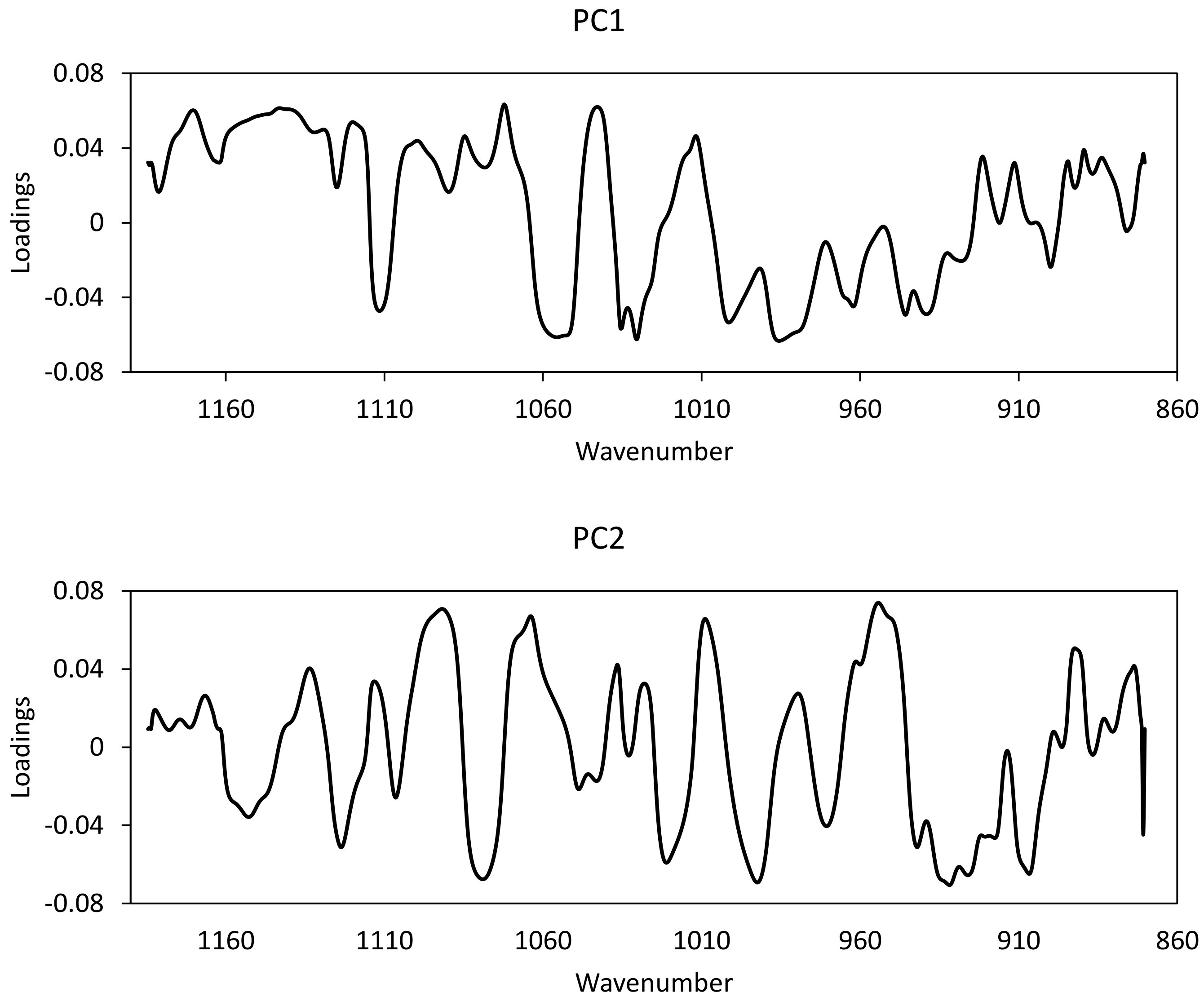


**Supplementary Figure S7.** Principal Component Analysis (PCA) loading profiles of FTIR cell wall polysaccharide spectra.
