## SUPPLEMENTARY FIGURES Supplementary Figure S1. Tissue-specific cellular responses and cross-sectional area dynamics across desiccation and rehydration for "Spatiotemporal pectin remodelling, glycoproteins, and LEA proteins maintain cell wall integrity during desiccation and rehydration in *Ramonda serbica*": Supporting_Information_File_S1.pdf

### Xtandem Parameter

Thierry Balliau

2026-09-01

#### Protein identifications

##### Xtandem Run information

| param | value |
|---|---|
| version | X! Tandem Alanine (2017.2.1.4) |
| sequence source #1 | /gorgone/pappso/moulon/users/thierry/20241018_vidovic_47/QC/contaminants_standarts.fasta |
| sequence source #2 | /gorgone/pappso/moulon/users/thierry/Projet/vidovic/database/total_ramonda.march2022.fasta |

##### spectrum filters

| param | value |
|---|---|
| neutral loss window | 0.02 |
| parent monoisotopic mass error minus | 15 |
| parent monoisotopic mass error plus | 25 |
| parent monoisotopic mass error units | ppm |
| maximum parent charge | 5 |
| fragment mass type | monoisotopic |
| fragment monoisotopic mass error | 15 |
| fragment monoisotopic mass error units | ppm |
| total peaks | 100 |
| mzFormat | 16 |
| neutral loss mass | 18.01057 |
| dynamic range | 100.0 |
| minimum fragment mz | 150.0 |
| minimum parent m+h | 500.0 |
| minimum peaks | 15 |
| timstof MS2 filters | chargeDeconvolution\|0.02dalton mzExclusion\|0.01dalton |
| parent monoisotopic mass isotope error | yes |

##### enzymatic cleavage

| param | value |
|---|---|
| cleavage site | [RK]\|{P} |
| cleavage semi | no |
| quick acetyl | yes |
| quick pyrolidone | yes |

| param | value |
|---|---|
| stP bias | yes |

##### Amino-acid modifications

| param | value |
|---|---|
| modification mass | 57.02146@C |
| potential modification mass | 15.99491@M, 15.99491@P |

##### refinment module

| param | value |
|---|---|
| use refine | yes |
| maximum missed cleavage sites | 0 |
| point mutations | no |
| maximum valid expectation value | 0.01 |
| modification mass | 57.02146@C |
| potential modification mass | 15.99491@M, 15.99491@W, 15.99491@P, 0.98402@N, 0.98402@Q |
| spectrum synthesis | yes |
| cleavage semi | no |
| NA | NA |
| use potential modifications for full refinement | yes |
| point mutations | no |
| unanticipated cleavage | no |

##### Score calculation

| param | value |
|---|---|
| minimum ion count | 4 |
| maximum missed cleavage sites | 1 |
| cyclic permutation | yes |
| include reverse | yes |
| b ions | yes |
| y ions | yes |
