## SUPPLEMENTARY FIGURES Supplementary Figure S1. Tissue-specific cellular responses and cross-sectional area dynamics across desiccation and rehydration for "Spatiotemporal pectin remodelling, glycoproteins, and LEA proteins maintain cell wall integrity during desiccation and rehydration in *Ramonda serbica*": Supporting_Information_script_S2.pdf

### template script label-free shotgun v.0.6.15

### This R script is designed to process and analyze data obtained by label-free shotgun proteomics.

#It can be used with the demo data sets available for download at https://entrepot.recherche.data.gouv.fr/dataset.xhtml?persistentId=doi:10.57745/9OKPZO.

### You should already have downloaded and installed R and MCQR on your computer (see https://forgemia.inra.fr/pappso/mcqr for instructions).

#-------------------------------------------------------- TABLE OF CONTENT ---------------------------------------------------------------


### 1. Setting up the R session

### 2. Analyzing quantification data obtained from extracted ion chromatogram (XIC)

###	2.1. Data and metadata loading

###	2.2. Quality checking

###	2.3. Data filtering

###	2.4. Intensity normalization

###	2.5. Post-normalization filters

###	2.6. Missing value imputation and aggregation of peptide intensities at the protein level

###	2.7. Descriptive analysis

###	2.8. Hypothesis testing

###	2.9. Exporting results


#****************************************************** 1. Setting up the R session

######### Load the MCQR package

```
library("MCQR")

packageVersion("MCQR")

	# [1] ‘0.6.15’
```

######### Create a directory for exporting results

```
path <- "./"

setwd (path)

newfolder <- "outmerged"

dir.create(file.path(path, newfolder))


#****************************************************   2. Analyzing quantification data
obtained from extracted ion chromatogram (XIC)


#########  2.1 Data and metadata loading


#----  Concatenate data and metadata

raw_xic_fCW <- mcq.read.masschroq(protFile="./parietal_proteins1.tsv",
pepFile="./peptides_q2_parietal1.tsv")

summary (raw_xic_fCW)

    # Number of proteins: 2812

    # Number of XICs: 152142

    # Number of peptides-mz: 9399

    # Number of peptides: 8690

    # Number of MS runs: 24

    # Peptide-mz intensities have not been normalized.

    # Peptide-mz intensities have not been imputed.

    # Number of NA values : 73434

    # % of NA values : 32.55


my_metadatam <- mcq.read.metadata("./metadata_final.ods")

summary(my_metadatam)

    # 4 levels for factor "condm" : bulk, DL+R1, HL+R3, R2

    # 8 levels for factor "repm" : 1, 2, 3, 4, 5, 6, 7, 8

    # 3 levels for factor "batch" : 3, 2, 1
```

```
        # 6 levels for factor "cond" : bulk, DL, HL, R1, R2, R3
        # 5 levels for factor "rep" : 1, 2, 3, 4, 5

    raw_xic <- mcq.add.metadata(raw_xic, metadata=my_metadatam)

#----  Retain only cell wall proteins
    good_accession <- read.table("./filtered_CW_proteins.csv", header=T, sep=",", dec=".",
stringsAsFactors=F)
    raw_xic <- mcq.select(raw_xic, factor="accession", levels=good_accession$accession)
    summary(raw_xic)
        # Number of proteins: 358
        # Number of XICs: 19568
        # Number of peptides-mz: 1221
        # Number of peptides: 1145
        # Number of MS runs: 24
        # Peptide-mz intensities have not been normalized.
        # Peptide-mz intensities have not been imputed.
        # Number of NA values : 9736
        # % of NA values : 33.22

#########  2.2. Quality checks (QC)

#----  Retention time-based QC
    mcq.plot.rt.alignment(directory="./time", ncol=6, nrow=4, ylim=c(-200, 200) ,
file="./outmerged/alignmentCW.pdf" )

    #-FiguresS3c,e,f
    mcq.plot.peak.width(raw_xic, file="./outmerged/peak_width_raw.pdf")
    mcq.plot.rt.variability(raw_xic, width=1, file="./outmerged/rt_var_raw.pdf")
```

```
mcq.plot.rt.variability(raw_xic, limit=100, width=1, file="./rt_var_zoom_raw.pdf")

mcq.plot.intensity.profile(raw_xic, factorToColor=c("cond"), interactiveView=TRUE)

mcq.plot.intensity.profile(raw_xic, factorToColor=c("cond"), file="./outmerged/profile_raw.pdf")


#----  Intensity-based QC

mcq.plot.intensity.violin(raw_xic, factorToColor=c("cond"), file="./outmerged/violin_raw.pdf")


#-FiguresS3d

mcq.plot.intensity.distribution(raw_xic, overlay=TRUE, file="./outmerged/intensity_distrib_raw.pdf")

mcq.plot.intensity.correlation(raw_xic, ref="msruna4", ncol=3, nrow=4, file="./outmerged/correlation_raw.pdf")

pca_on_raw_xic = mcq.compute.pca(raw_xic)

mcq.plot.pca(pca_on_raw_xic, factorToColor=c("cond"), tagType="label", labels=c("cond","rep"), file="./outmerged/pca_raw_cond.pdf" )

mcq.plot.pca(pca_on_raw_xic, factorToColor=c("batch"), tagType="label", labels= c("cond", "rep", "batch"), file="./outmerged/pca_raw_batch.pdf" )

mcq.plot.pca(pca_on_raw_xic, factorToColor=c("condm"), tagType="label", labels=c("condm","repm"), file="./outmerged/pca_raw_condm.pdf" )


#----  Chromatographic peaks based-QC

mcq.plot.counts(raw_xic, file="./outmerged/counts_raw.pdf")


#########  2.3. Data filtering


filtered_xic <- raw_xic


#----  Filter peptide-mz showing very large chromatographic peaks

filtered_xic <- mcq.drop.wide.peaks(filtered_xic, cutoff=200)
```

#---- Filter peptide-mz showing very highly variable retention time

filtered_xic = mcq.drop.variable.rt(filtered_xic, cutoff=20)

#---- Filter peptide-mz showing at the end of the chromatographic run

filtered_xic= mcq.drop.rt.range(filtered_xic, rtmin=0, rtmax=875)

#---- Checking the effects of the filters

mcq.plot.peak.width(filtered_xic, file="./outmerged/peak_width_filtered.pdf")

mcq.plot.rt.variability(filtered_xic, width=1, file="./outmerged/rt_var_filtered.pdf")

mcq.plot.intensity.profile(filtered_xic, factorToColor=c("cond"), file="./outmerged/profile_filtered.pdf")

mcq.plot.intensity.profile(filtered_xic, factorToColor=c("condm"), file="./outmerged/profile_filteredm.pdf")

######### 2.4. Intensity normalization

normalized_xic=mcq.compute.normalization(filtered_xic, method="median")

#---- Checking the effects of normalization (FigureS3a)

mcq.plot.intensity.violin(normalized_xic, factorToColor=c("cond"), file="./outmerged/violin_method21.pdf")

pca_on_normalized_data = mcq.compute.pca(normalized_xic)

mcq.plot.pca(pca_on_normalized_data, factorToColor=c("condm"), tagType="label", labels= c("condm","repm"), file="./outmerged/pca_filtered_method21.pdf" )

mcq.plot.intensity.profile(method2, factorToColor=c("condm"), file="./outmerged/profile_norm_method21.pdf")

######### 2.5. Post-normalization filters

#---- bulk removal

normalized_xic_nobulkpost <- mcq.drop(normalized_xic, factor="condm", level=c("bulk"))

#---- Shared peptides removal

mcq.plot.peptide.sharing(normalized_xic_nobulkpost, file="./outmerged/shared_nobulkpost.pdf")

normalized_xic_nobulkpost = mcq.drop.shared.peptides(normalized_xic_nobulkpost)

#---- Remove peptide-mz with too many missing values

mcq.plot.peptide.reproducibility(normalized_xic_nobulkpost, file="./outmerged/repro_nobulkpost.pdf")

normalized_xic_nobulkpost = mcq.drop.unreproducible.peptides(normalized_xic_nobulkpost, method="refined", percentNA=0.25, flist="cond")

#---- Remove peptide-mz which are not correlated to the peptide-mz belonging to the same protein

normalized_xic_nobulkpost <- mcq.drop.uncorrelated.peptides(normalized_xic_nobulkpost, rCutoff=0.5)

#---- Remove proteins quantified with less than 2 peptides

normalized_xic_nobulkpost <- mcq.drop.proteins.with.few.peptides(normalized_xic_nobulkpost, npep=2)

summary(normalized_xic_nobulkpost)

### Number of proteins: 43

### Number of XICs: 2082

### Number of peptides-mz: 112

### Number of peptides: 112

### Number of MS runs: 19

### Peptide-mz intensities have been normalized.

### Peptide-mz intensities have not been imputed.

### Number of NA values : 46

### % of NA values : 2.16

######### 2.6. Missing value imputation and peptide-to-protein aggregation

#---- Peptide-level imputation

```
knn_imputed_xic_nobulkpost <- mcq.compute.peptide.imputation(normalized_xic_nobulkpost, method = "knn")
```

#---- Peptide-to-protein aggregation

```
prot_abund_knn_nobulkpost <- mcq.compute.protein.abundance(knn_imputed_xic_nobulkpost, method="sum")
```

#---- Protein-level imputation

```
imputed_prot_abund_knn_nobulkpost <- mcq.compute.protein.imputation(prot_abund_knn_nobulkpost)
```

######### 2.7. Descriptive analysis of the protein data set

#---- PCA representation

```
pca_on_proteins_knn_nobulkpost <- mcq.compute.pca(imputed_prot_abund_knn_nobulkpost)

mcq.plot.pca(pca_on_proteins_knn_nobulkpost, factorToColor=c("condm"), labels =c("condm", "repm", "batch"), tagType="label", labelSize=3, file="./outmerged/pca_prot_knn_nobulkpost.pdf")

#-Figure 4b

mcq.plot.pca(pca_on_proteins_knn_nobulkpost, factorToColor=c("condm"), labels =c("condm"), tagType="label", labelSize=3, file="./outmerged/pca_prot_knn_nobulkpost1.pdf")

mcq.plot.pca(pca_on_proteins_knn_nobulkpost, factorToColor=c("condm"), labels =c("condm", "repm"), tagType="label", labelSize=3, file="./outmerged/pca_prot_knn_nobulkpost2.pdf")

#-Export PCA results

pca_on_proteins_results_knn_nobulkpost <- mcq.get(pca_on_proteins_knn_nobulkpost)

write.csv(pca_on_proteins_results_knn_nobulkpost$tca, file = "./outmerged/pca_samples_tca_nobulkpost.csv", row.names = FALSE)

write.csv(pca_on_proteins_results_knn_nobulkpost$tco, file = "./outmerged/pca_proteins_tco_nobulkpost.csv", row.names = FALSE)
```

```
#----  Heatmap representation

        mcq.plot.heatmap(imputed_prot_abund_knn_nobulkpost, flist=c("condm"), factorToColor=c("condm"), protLab=FALSE, file="./outmerged/heatplot_knn_nobulkpost.pdf")

        #-Figure S4

        mcq.plot.heatmap(imputed_prot_abund_knn_nobulkpost, flist=c("condm", "repm"), factorToColor=c("condm"), protLab=FALSE, file="./outmerged/heatplot_knn_nobulkreppost.pdf")


#----  CV distribution

        cv_prot_knn_nobulkpost <- mcq.compute.cv(imputed_prot_abund_knn_nobulkpost)

        #-FigureS3b

        mcq.plot.cv(cv_prot_knn_nobulkpost,file="./outmerged/cv_knn_nobulkpost.pdf" )

        cv_by_cond <- mcq.compute.cv(imputed_prot_abund_knn_nobulkpost, flist = "condm")


        #-in which conditions are the most variable proteins?

        df_cv_cond <- mcq.get.dataframe(cv_by_cond)

        aggregate(cv ~ condm, data = df_cv_cond, FUN = mean)

        high_cv_cond <- df_cv_cond[df_cv_cond$cv >= 70, ]

        table(high_cv_cond$condm)

        #DL+R1 HL+R3    R2

        #7     4     4


        #-Figure 4b

        df_cv <- cv_prot_knn_nobulkpost@resultcv

        if(!is.data.frame(df_cv)){df_cv <- data.frame(cv = as.numeric(df_cv))}

        file_path <- "./outmerged/CV_Sirovipost.csv"

        write.csv(df_cv, file = file_path, row.names = FALSE)


#----  Protein clustering
```

```
protein_clusters_knn_nobulkpost <- mcq.compute.cluster(imputed_prot_abund_knn_nobulkpost, flist=c("condm"), nbclust=c(2,6), method=c("kmeans"))

        # Clustering Methods:
        #   kmeans
        # # Cluster sizes:
        #   6
        # # Validation Measures:
        #   6
        # # kmeans APN          0.1683
        # AD           0.4340
        # ADM          0.2281
        # FOM          0.2399
        # Connectivity  21.5813
        # Dunn         0.0954
        # Silhouette    0.5662
        # # Optimal Scores:
        #   #   Score   Method Clusters
        # APN         0.1683 kmeans 6
        # AD          0.4340 kmeans 6
        # ADM         0.2281 kmeans 6
        # FOM         0.2399 kmeans 6
        # Connectivity 21.5813 kmeans 6
        # Dunn        0.0954 kmeans 6
        # Silhouette   0.5662 kmeans 6

#-FigureS5

        mcq.plot.cluster(protein_clusters_knn_nobulkpost, method="kmeans", nbclust=6, file="./outmerged/cluster_knn_nobulkpost.pdf")
```

my_clustering_results_knn_nobulkpost <- mcq.compute.cluster(imputed_prot_abund_knn_nobulkpost, flist=c("condm"), nbclust=c(6), method=c("kmeans"))

my_clustering_results_knn_nobulkpost<- mcq.get(my_clustering_results_knn_nobulkpost)

#-TableS4

write.csv(  my_clustering_results_knn_nobulkpost,file = "./outmerged/xic_clustering_results_knn_nobulkpost.csv", row.names = FALSE)

######### 2.8. Hypothesis testing

#----  Remove proteins with maximum fold change < 1.5

imputed_filtered_prot_abund_knn_nobulkpost <- mcq.drop.low.fold.changes(imputed_prot_abund_knn_nobulkpost, cutoff=1.5, flist=c("condm"))

#----  ANOVA

anova_on_proteins_knn_nobulkpost <- mcq.compute.anova(imputed_filtered_prot_abund_knn_nobulkpost, flist=c("condm"), inter=T)

mcq.plot.pvalues(anova_on_proteins_knn_nobulkpost, file="./outmerged/pvalues_knn_nobulkpost.pdf")

#----  Post-hoc Tukey test

proteins_signif_cond_knn_nobulkpost <- mcq.select.pvalues(anova_on_proteins_knn_nobulkpost, padjust=TRUE, 0.05, flist="condm"))

tukey_cond_knn_nobulkpost <- mcq.compute.tukey(anova_on_proteins_knn_nobulkpost, flist="condm", protlist=proteins_signif_cond_knn_nobulkpost)

mcq.plot.tukey(tukey_cond_knn_nobulkpost, qprot=imputed_filtered_prot_abund_knn_nobulkpost, factorToColor="condm", file="./outmerged/tukey_cond_knn_nobulkpost.pdf")

######### 2.9. Extracting and exporting analysis results

#---- Extracting and exporting protein quantification data

protein_abundance_table_knn_nobulkpost <- mcq.get(imputed_filtered_prot_abund_knn_nobulkpost)

write.csv( protein_abundance_table_knn_nobulkpost, file = "./outmerged/protein_abundance_table_knn_nobulkpost.csv", row.names = TRUE)

#---- Extracting and exporting ANOVA results

#-table S5.2

my_anova_results_knn_nobulkpost <- mcq.get(anova_on_proteins_knn_nobulkpost)

write.csv(my_anova_results_knn_nobulkpost,file = "./outmerged/anova_results_knn_nobulkpost.csv", row.names = FALSE)

#---- Extracting and exporting a table merging ANOVA results with protein quantification data

#-table S5.1

merged_datapost <- merge(protein_abundance_table_knn_nobulkpost,my_anova_results_knn_nobulkpost, by = "accession",all.x = TRUE)

write.csv( merged_datapost,file = "./outmerged/merged_results_knn_nobulk1post.csv", row.names = FALSE)
